# Cross-species Transcriptomic Network Analysis Reveals Links Between Ribosomal Protein Mutation and Cancer

**DOI:** 10.1101/2020.12.11.420604

**Authors:** Jai A Denton, Mariana Velasque, Floyd A Reed

## Abstract

Ribosomal proteins (RPs) are critical to all cellular operations through their key roles in ribosome biogenesis and translation, as well as their extra-ribosomal functions. Leveraging previously identified RP mutants we characterised the RP macro transcriptome and sought to compare it with transcriptomes of pathologies associated with failures of ribosomal function, cancer and Zika virus (ZIKV). Single gene-based analysis revealed highly variable transcriptomes of RP mutations with little overlap in genes that were differentially expressed. In contrast, weighted gene co-expression network analysis revealed a highly conserved transcriptomic network pattern across all RP mutants studied. In addition, when we compared network changes in RP mutants, we observed similarities to transcriptome alterations in human cancer, and thus confirming the oncogenic role of RPs. Finally, it is known that ZIKV infection influences translational machinery, but this study shows infection network changes dissimilar to those of either the RP mutation or cancer.

## Introduction

Gene networks are typically more than the sum of their parts. As new components are added to a network, complexity can increase exponentially, such that it is difficult to predict or even assess the reverberations the consequences small changes have throughout entire networks (Crow et al., 2019). Therefore when examining changes brought on by the modification of a single network component, it is important to examine not just how other single components are affected but how networks themselves change. As networks grow in size and influence a greater number of cellular processes, this becomes more critical.

Ribosomal proteins (RPs) are a diverse group of highly conserved proteins that are central to ribosome biogenesis. Given the dependency of normal cellular function on rapid and accurate protein production, it is unsurprising that mutations in RP-encoding genes are highly deleterious, resulting in a variety of adverse outcomes (Wang et al., 2015; Yassin et al., 2005). Thus, homozygous deletion or mutation of RPs is typically lethal (McGowan et al., 2008), whereas heterozygous RP mutations are typically haploinsufficient with a fitness cost (Kim et al., 2010; Marygold et al., 2007; Weijers et al., 2001; Zheng et al., 2016). In addition to their role in ribosome biogenesis and function, some RPs have important extra-ribosomal activities (Warner and McIntosh, 2009). Their mis-expression has been detailed in numerous aberrant cellular processes. In recent years, RPs have been shown to participate in stress responses, nucleolar integrity, cell cycle control, cell proliferation, genome integrity, telomere length and cell death (Abdulkina et al., 2019; Lai et al., 2009; Moin et al., 2016; Nicolas et al., 2016). Thus, ribosome biogenesis is highly dose-dependent, with small changes impacting numerous cellular processes.

Due to this diverse array of cellular functions, RP mutations potentiate cancer. Suboptimal ribosome biogenesis, or incorrect function of RPs generally result in oncogenesis (Lai et al., 2009; Oršolić et al., 2020), whereas their normal functioning is tumor-suppressive (Amsterdam et al., 2004; Fancello et al., 2017). As such, RP mutations are associated with diverse cancers (De Keersmaecker et al., 2013; Rao et al., 2012). Moreover, hyperactivation of ribosome biogenesis is also frequently seen in cancer cells (Dolezal et al., 2018). Impaired ribosome biogenesis arrests cell cycle progression in a p53-dependent manner (Volarević et al., 2000; Pestov et al., 2001). This occurs through RP-mediated stabilization of p53, with critical involvement of RpL5 and RpL11 (Bursać et al., 2012). Mutant forms of these two RPs have been documented in many cancers (Dong et al., 2017). In addition, RpL22, RpL10, RpS15, RpS20 and RpL5 are also mutated in diverse human cancers. Moreover, patients with Diamond-Blackfan anemia, which results from mutations in a wide range of RPs, are at a higher risk of developing cancer (Boria et al., 2010). Thus, either directly or indirectly, by virtue of being highly pleiotropic in cellular processes, RPs are critical in oncogenesis.

RPs in *Drosophila melanogaster* predominantly include a class of mutants known as the Minute loci (Bridges et al., 1923; Kongsuwan et al., 1985; Lambertsson, 1998; Marygold et al., 2007). First described in *D. melanogaster* almost a century ago (Bridges et al., 1923), the Minute loci were originally named for their resulting smaller bristles. In addition to thinner bristles, Minutes bearing flies exhibit delayed development, lower viability and fertility, and altered body size (Bridges et al., 1923; Marygold et al., 2007). These are dominant phenotypes due to haploinsufficiency of RPs, and with rare exception, are homozygous lethal (Marygold et al., 2007).

Despite comprehensive study of RPs and Minute loci in *D. melanogaster*, regulatory response and cellular outcomes remain little understood. For instance, despite the pleiotropic nature of RPs, it is unknown whether disruption of RPs causes a standard cellular or a transcriptomic response. Recent work identified XRP1, a DNA-binding protein involved in genome stability, as essential to regulate the cellular response to aberrant RpS3 and RpS17 expression in *D. melanogaster* (Lee et al., 2018). It was also shown that mutations in the RpS3 or mahjong genes activate Toll and oxidative stress pathways and the Nrf2 (cnc) stress factor (Kucinski et al., 2017). Given the pleiotropic and variable nature of RP disruptions, further analysis is required to determine whether RPs share a transcriptomic response or if these mutations shape the transcriptome on an individual basis.

In the present study, we have investigated whether there are consistent single gene and network transcriptomic responses to RP mutations in *D. melanogaster* using whole-body RNA sequencing data from 7 RP mutant lines. We have selected Minute loci arising from *P*-element disruption of RP-encoding genes, identified on the basis of their phenotypes and the location of *P*-element genomic insertions. Furthermore, we sought to contextualise transcriptomic responses using available transcriptomic data from cancer cells.

## Results

### Genomic Locations of Ribosomal Protein-Encoding Genes

Unlike many gene families RPs are not clustered within the genome. If the distribution of RPs in the genome were random, it should result in exponentially distributed distances between adjacent RPs. However, we found that their distribution was consistent with non-clustered, essentially random placement in the genome (n=86, KS-test D=0.1121, P=0.2134) (Figure 1).

**Figure 1.**
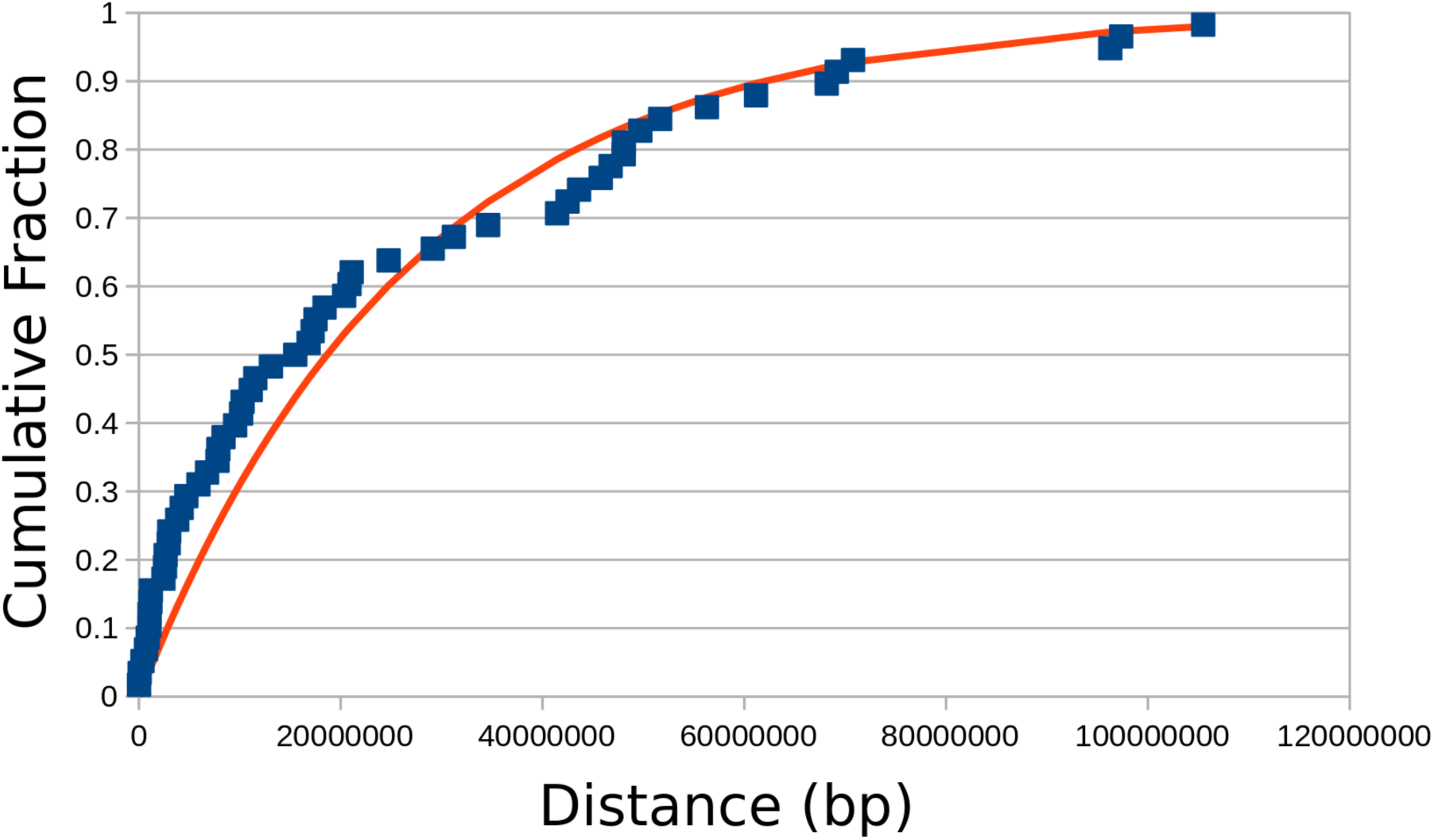
RPs are distributed essentially randomly in the *Drosophila* genome. The cumulative fraction shown in blue and the theoretical exponential curve based on a Poisson process is plotted for comparison in red. RP placement is random (n=86, KS-test D=0.1121, P=0.2134).

### Aberrant Expression of RPs

RP mutants were generated using *P*-element mutagenesis and have been previously confirmed as Minutes via phenotypic screens (Marygold et al., 2007). We detected aberrant expression of target RPs in five of seven Minute *P*-element mutagenesis lines used when compared to an unmutated control (adj-p<0.05) (Table 1). RNA sequencing to identify transcriptomic changes was successfully performed on single flies from seven RP mutagenesis lines (Materials & Methods). With the exception of RpS19b, each line displays reduced expression of the target RP, ranging from −0.03 to −0.91-fold (log_2_)change (Table 1). However, RpS19b was upregulated almost 5-fold (log_2_). As the P-element insertion for the RpS19b mutant is 34 bp upstream from the start codon, the inserted promoter is likely driving the over expression (FlyBase - FBrf0230790/FBti0181318). Although previous research confirmed the Minute phenotype and characterised the genome insertion (Marygold et al., 2007), our analysis did not detect a significant change in the target RPs, RpL3 or RpL30 (adj-p > 0.05). These were also the two smallest expression changes, 0.03 and 0.17, respectively. However, despite the absence of a statistically significant reduction in expression, RpS19b, RpL3, and RpL30 are included in the analysis as they are confirmed mutants with characterised phenotypes.

**Table 1.**
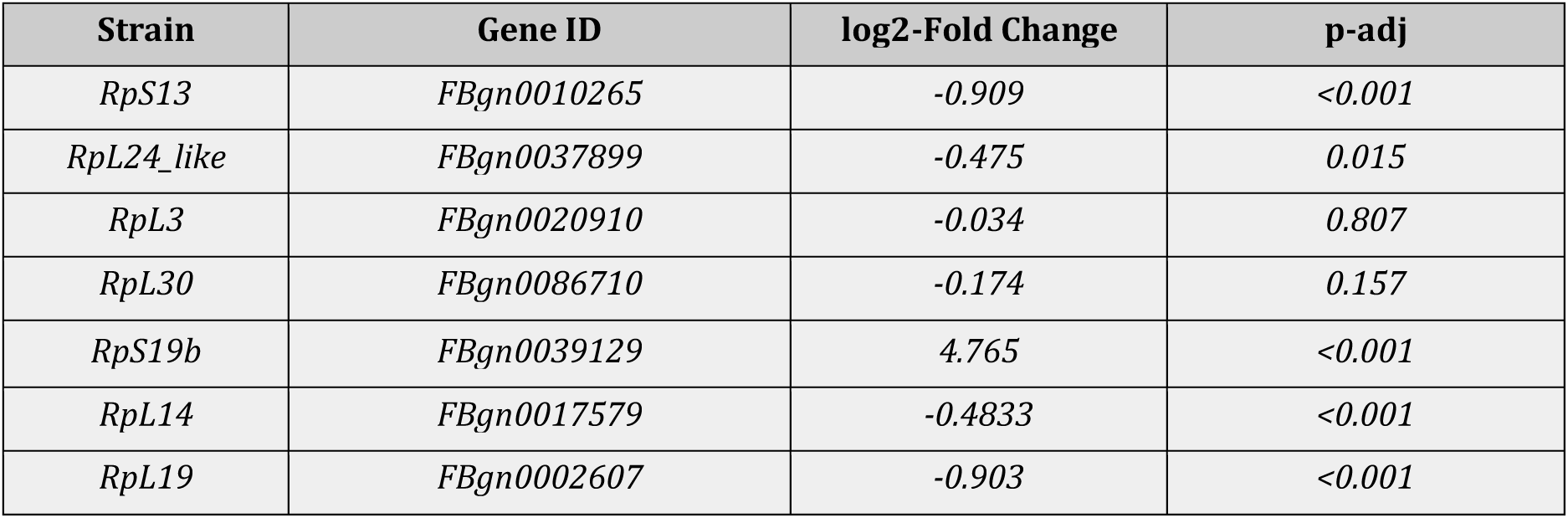
Ribosomal Protein Gene Expression. Expression values for each of the affected ribosomal protein-encoding genes in their respective mutant backgrounds compared to an control. For example, Strain RpL14 contains expression data for RpL14. Each transcript had comparatively lower expression, consistent with a knockdown, except RpS19, which had considerably higher expression. Neither RpL3 nor RpL30 had significantly different expression (adj-p > 0.05). The strain names refers to gene affected but is only a single allele (Table 7)

| Strain | Gene ID | log2-Fold Change | p-adj |
| --- | --- | --- | --- |
| *RpS13* | *FBgn0010265* | -0.909 | <0.001 |
| *RpL24_like* | *FBgn0037899* | -0.475 | 0.015 |
| *RpL3* | *FBgn0020910* | -0.034 | 0.807 |
| *RpL30* | *FBgn0086710* | -0.174 | 0.157 |
| *RpS19b* | *FBgn0039129* | 4.765 | <0.001 |
| *RpL14* | *FBgn0017579* | -0.4833 | <0.001 |
| *RpL19* | *FBgn0002607* | -0.903 | <0.001 |

### The Minute Transcriptome

There is no strong shared transcriptomic response between Minute mutations. Each of the Minute strains tested displayed a considerable number of genes that were differentially expressed relative to a control strain (adj-p <0.05) (Supplementary Figure 1, Table 2). The data show large numbers of differentially expressed genes, typically with small magnitude changes. There is also considerable variation in the number of differentially expressed genes, with RpL24-like having 3350 and RpS13 having 469. As such there is unlikely to be a clearly defined core of genes among RP mutants.

**Table 2.**
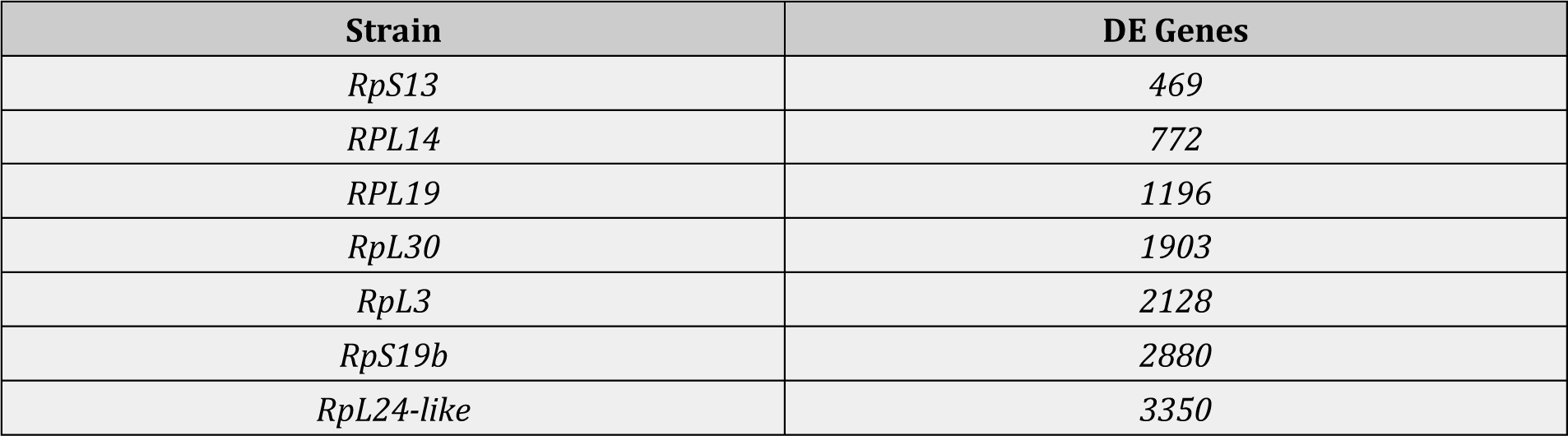
Differentially expressed genes. The number of differentially expressed (DE) genes in each treatment (strain). The strain names refers to gene affected but is only a single allele (Table 7)

There are only 18 genes differentially expressed across all RP mutants (Table S1). Even with the removal of RpS13, which raises shared differentially expressed genes to 67, a clearly defined transcriptomic response is lacking. In the absence of a single core transcriptome, we explored the possibility of overlapping groups of differentially expressed genes. This analysis shows considerable overlap between various groups (Figure 2). For example, RpS30, RpS3, RpS19b and RpL24-like share 441 differentially expressed genes.

**Figure 2.**
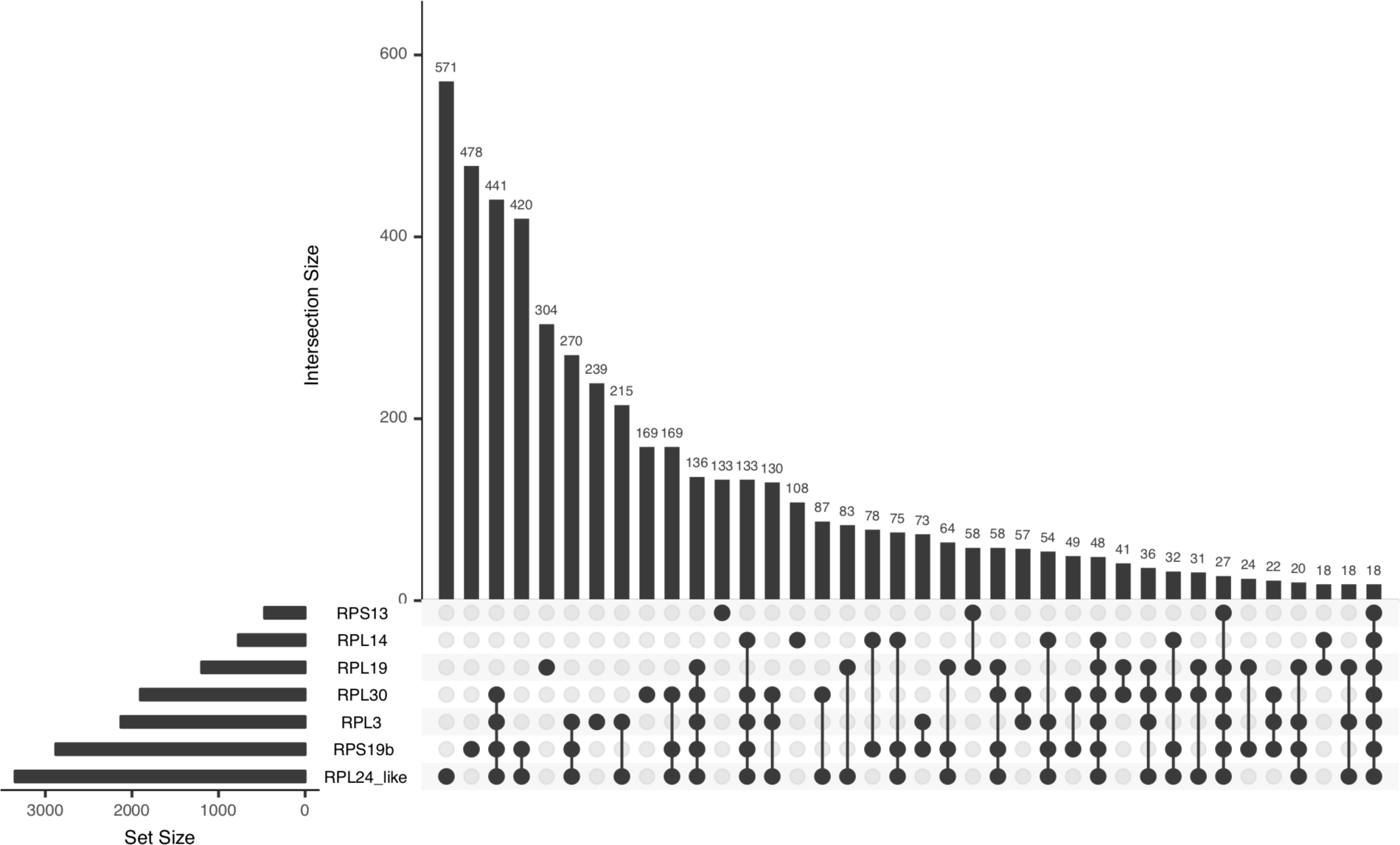
UpSetR diagram detailing overlapping differentially expressed genes between ribosomal protein (RP) mutants. A Venn diagram-like representation of differentially expressed (DE) genes exhibiting statistically significant changes (adj p-value <0.05). Set Size denotes the total number of DE genes. Linked dots and corresponding column indicate the number of DE genes shared between those RP mutants. Single dots indicate the number of DE genes unique to the RP mutant.

### Single-Gene Misregulation

As no clear pattern of expression was observed across all mutants in response to mutations in RP-encoding genes, both previously identified regulatory genes and markers for implicated pathways were examined.

Previous work identified XRP1 and IBRBP1 as being upregulated in response to aberrant RP expression (Lee et al., 2018; Blanco et al., 2020). However, we were unable to identify XRP1 or IBRBP1 as a clear marker. Only in the RpS13 line was XRP1 upregulated (adj p-value < 0.05) (Table 3). IBRPB1 was not upregulated in any line, although it was downregulated in RpL24-like (Table 3).

**Table 3.** XRP1 / IRBP1 Expression in each RP mutant strain.

|  | XRP1 |  | IBRBP1 |  |
| --- | --- | --- | --- | --- |
|  | log2 - Fold Change | p-adj | log2-Fold Change | p-adj |
| RpS13 | <u>0.648</u> | <u>0.0346</u> | 0.223 | 0.416 |
| RpL24-like | -0.1609 | 0.5723 | <u>-0.364</u> | <u>0.038</u> |
| RPL3 | -0.1816 | 0.5448 | -0.152 | 0.446 |
| RPL30 | -0.2951 | 0.2863 | -0.150 | 0.447 |
| RpS19b | -0.3835 | 0.1371 | -0.275 | 0.124 |
| RpL14 | -0.453 | 0.1181 | -0.100 | 0.656 |
| RpL19 | 0.0911 | 0.8155 | -0.012 | 0.969 |

As several pathways controlled by a handful of key genes have also been implicated in the cellular response to RP mutations, a candidate gene-based approach was undertaken. Of the 20 candidate genes examined, only asp, CycB3, pik92e, ask1 & bsk were differentially expressed in four or more of the RP mutants (Supplementary Table S2).

### Minute Gene Ontology

Although there are very few differentially regulated core genes, there is a large core of gene ontology (GO) categories shared among all seven mutant lines. We classified differentially expressed genes for each of the mutant lines based on biological function GO. 138 GO categories were shared by all mutant lines (Supplementary Table S3). These categories included RNA processing, translation, and programmed cell death. Moreover, using pairwise comparisons, it is clear that there is considerable overlap in the biological processes affected by RP mutations (Figure 3). However, GO analysis based on differentially expressed genes can be limited especially when large numbers of genes are differentially expressed (Boyle et al., 2017). Thus we employed network analysis to provide a more accurate view of how the transcriptome is shaped by RP mutations.

**Figure 3.**
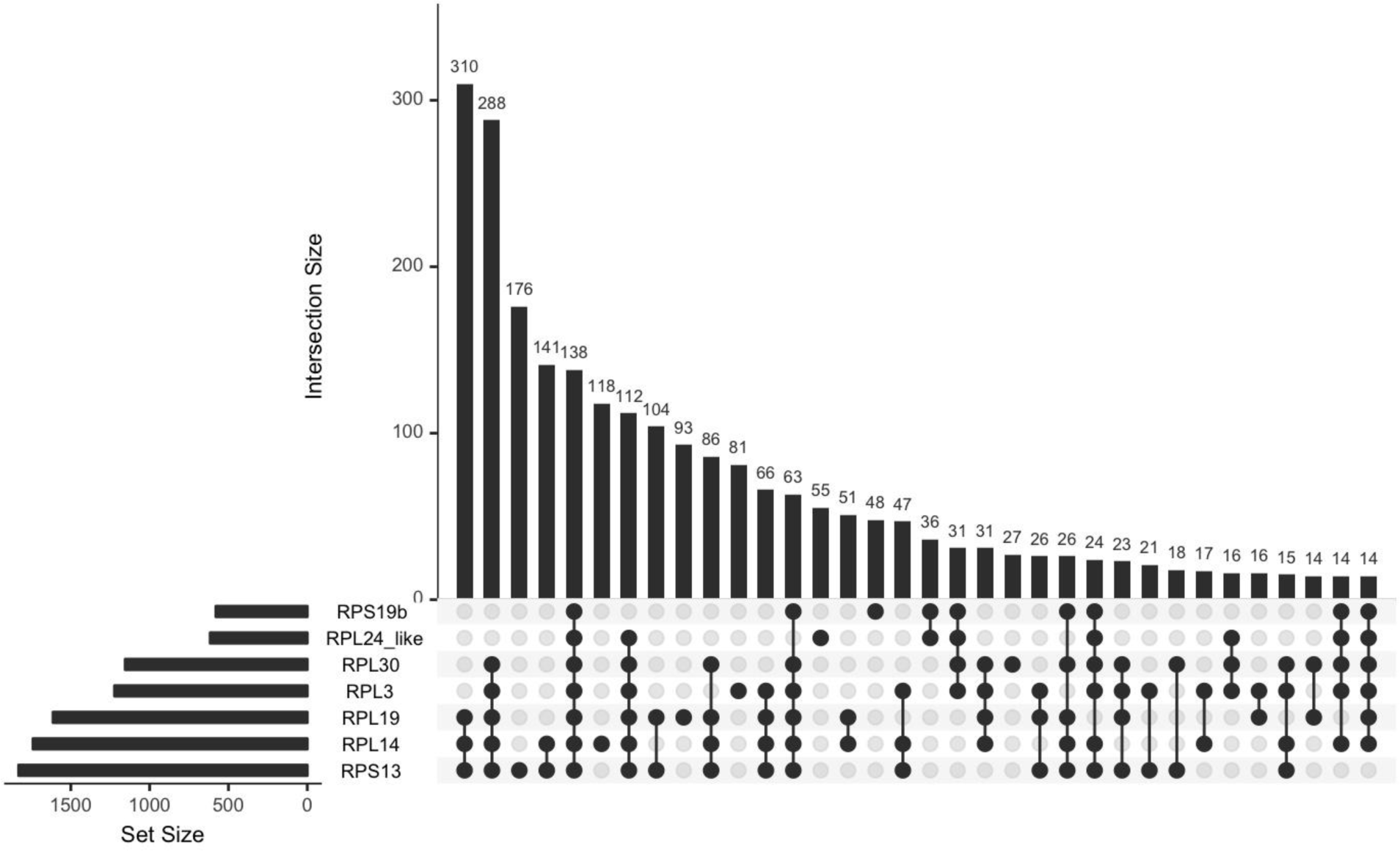
UpSetR diagram detailing overlapping biological process gene ontology (GO) categories of differentially expressed genes in each of the RP mutants. A Venn diagram-like representation of GO categories for the differentially expressed (DE) genes (adj p-value <0.05). Set Size denotes the total number of GO categories. Linked dots and the corresponding column indicate the number of GO categories shared between those RP mutants. Single dots indicate the number of GO categories unique to that specific RP mutant.

### RP Network Analysis

Genes are not independent. Their activation and expression depend on a series of interactions with other genes and pathways. Genes with similar biological functions tend to be regulated by the same transcription factor and, overall, have similar expression profiles (Weirauch, 2011). Therefore, associations can be identified by clustering groups of genes with similar expression (i.e. co-regulated) into modules, improving functional genome annotation using the *guilt-by-association principle* (Wolfe et al., 2005). Similarly, differences in network co-expression patterns between two groups or treatments can be used to indicate gene clusters involved in a particular pathway. Differential network analysis, where differences in gene co-expression between two data sets can be used to identify condition-associated pathways, leverages these co-expression patterns and is an alternative to typical differential gene expression analysis (Bhuva et al., 2019; Hsiao et al., 2016).

We first explore the differences between each RP mutant strain and the control and identify structural changes in pathways associated with ribosome biogenesis dysregulation (RP Differential Network Analysis). Although knowledge of changes in ribosomal pathway in Minutes are crucial to understand how ribosomes are regulated, the information on the whole cluster (differentially or not differentially co-expressed) is missing. Therefore, we also attempted to identify the ribosomal hub modules by identifying clusters of genes co-expressed in each Minute strain (RP Weighted Gene Co-expression Network Analysis).

#### RP Differential Network Analysis

The Differential Network Analysis (Materials & Methods) was used to identify differences in pathways between Minutes and control were tested via permutation. A total of 6004 unique genes were found to be differentially co-expressed in at least one RP mutant strain (Table 4). The data show a large number of differentially co-expressed (DCE) genes within the ribosomal pathway. Although all networks have shown high connectivity between DCE genes, there is a large variance in magnitude across the DCE genes and large core of shared genes (61). Similar to the DE analysis, we were unable to identify XRP1 or IBRBP1 as part of the ribosomal network.

**Table 4.** Differentially co-expressed genes. The number of differentially co-expressed (DCE) genes in each treatment (strain) pathway. The strain names refers to gene affected but is only a single allele (Table 7)

| Strain / Gene | DCE Genes |
| --- | --- |
| RpL3 | 2640 |
| RpL14 | 1426 |
| RpS13 | 1143 |
| RpS19b | 3203 |
| RpL19 | 1874 |
| RpL24-like | 2580 |
| RpL30 | 3119 |

Although we identified this 61 gene core, no significant gene ontology categories were shared between all strains. Differentially co-expressed genes were classified based on the on biological function GO and a total of 171 GO (Figure 4) categories were present in at least one of the mutant lines (Supplemental material S2)

**Figure 4.**
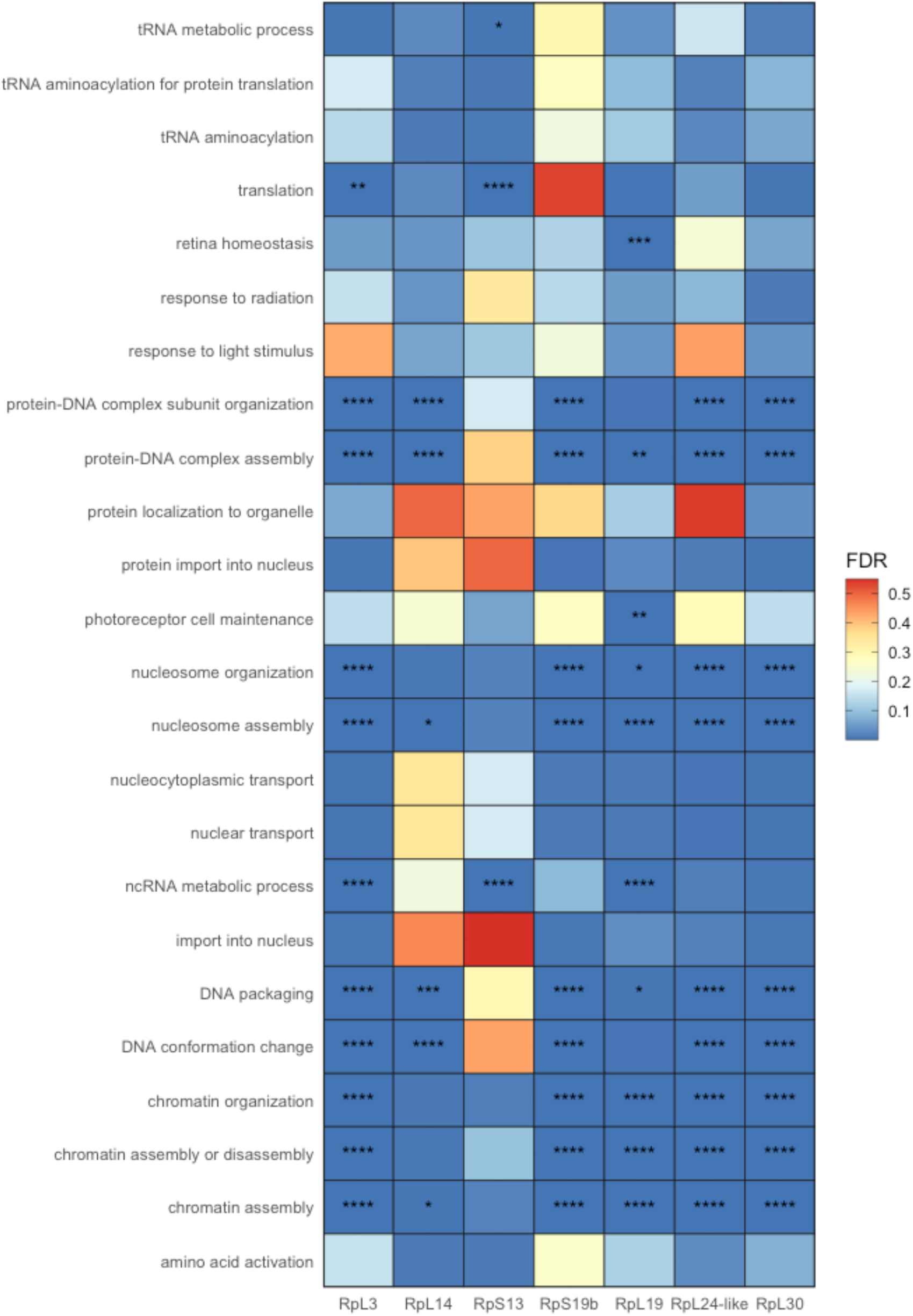
List of Gene Ontology (GO) terms that were significantly enriched among differentially co-expressed genes in RP mutant strains. The gradient (red to blue) shows the false discovery rate for each GO term. Asterisks denote statistically significant enrichment (p < 0.05), double, triple and four asterisks denote significant enrichment (p < 0.01, p < 0.001 and p < 0.0001 respectively).

### RP Weighted Gene Co-expression Network Analysis

The weighted gene co-expression network analysis (WGCNA) (Materials & Methods) is a systematic biological method used to describe patterns of gene associations between samples and treatment groups. It uses expression of thousands of genes to construct co-expression modules related to a phenotype or treatment group. Thus, it has been extensively used in genome studies to identify gene sets of interest or gene networks (Langfelder and Horvath, 2008).

We attempted to understand how changes in RP expression affects modules through changes in gene clustering. In other words, we investigated whether RP expression of mutants had a different clustering pattern when compared to controls (Spradling et al., 1999). The transcriptomic network of each RP mutant was contrasted with that of the wild type. With network construction, we measured intramodular gene connectivity between mutants, selecting highly connected genes, defined as RP hub genes. These hub genes were highly connected in their respective RP modules and are likely to serve critical functions in regulating ribosomal activity. Thus, for each mutant, we identified hub genes and their patterns of expression, identifying a list of differentially expressed genes (see S2 Supplemental Materials for a full list). We also investigated whether the expression of such RP hub genes was conserved across experimental groups.

Module preservation is a measure of how gene clusters (small subsets within a network) from two different sets of data have similar expression profiles. Preservation is measured using Zsummary results and respective p-values. A Zsummary statistic of >10 is considered strongly similar, >5 moderately similar and <2 as dissimilar. Our Zsummary scores (>10) indicate stronger evidence that a module is well preserved across all RP mutants (Table 5). p-values for WGCNA Module preservation analysis ranged between 5.89E-12, for RpL3 / RpS13, to 1.04E-78, for RpS19b / RPL30. This suggests a well conserved transcriptomic response among all RP mutants. For example, within the RpL3 mutant transcriptome, the module containing all genes that grouped with RpL3 was compared to the module grouping all RpL14-responsive genes from the RpL14 mutant.

**Table 5.** WGCNA Module Preservation. The reported Zsummary, shown in blue, and p-values, shown in yellow, of module correlations between each mutant strain. This is a pairwise comparison between a pair of modules containing the strain-specific RP mutation (e.g., *RpL3*-containing module in the RpL3 strain compared against the *RpL14*-containing module in the RpL14 strain).

|  | RpL3 | RpL14 | RpS13 | RpS19b | RpL19 | RpL24-like | RpL30 |
| --- | --- | --- | --- | --- | --- | --- | --- |
| RpL3 |  | <0.0001 | <0.0001 | <0.0001 | <0.0001 | <0.0001 | <0.0001 |
| **Rpl14** | 12.6465 |  | <0.0001 | <0.0001 | <0.0001 | <0.0001 | <0.0001 |
| **RpS13** | 13.3311 | 17.1543 |  | <0.0001 | <0.0001 | <0.0001 | <0.0001 |
| **Rp19b** | 21.5842 | 16.7581 | 10.8456 |  | <0.0001 | <0.0001 | <0.0001 |
| **RpL19** | 17.4687 | 17.5810 | 14.9769 | 19.5649 |  | <0.0001 | <0.0001 |
| **RpL24-like** | 22.1592 | 15.0877 | 11.2195 | 24.7626 | 16.9910 |  | <0.0001 |
| **RpL30** | 22.0449 | 16.4723 | 13.3747 | 26.2302 | 20.2664 | 24.6987 |  |

Network analysis not only showed that components of target modules were conserved, but highlighted the preserved nature of the RP gene network, suggesting a common activation pathway (see S2 Supplemental Materials for a full list of shared genes). Using GO analysis, we also investigated biological processes of RP hub genes (Supplementary Figures 1-7). However, we did not find any significantly enriched group in our analysis. Furthermore, the module preservation analysis also revealed that the majority of genes present on the network are shared by multiple RP modules.

The high degree of module preservation between RP mutant strains means that direct comparisons can be conducted between these strains and other transcriptomic datasets with proposed RP-driven effects. This allows us to seperate transcriptomes that are a product of RP-based changes from those wherein RPs are affected as a result.

### RPs, Cancer & Zika Virus - Comparative WGCNA

As RPs are highly conserved across the Eukaryota and glioblastoma and breast cancer have been previously attributed to disruption of RP genes (Penzo et al., 2019), we compared preservation levels between RP hub genes and their human counterparts. Moreover, although aberrant RP activity has been implicated in Zika virus infections, it has been proposed the aberrant activity is a response to rather than a driver of changes in the tissue of Zika virus infected individuals (Hetman and Slomnicki, 2019; Slomnicki et al., 2017). Therefore, cancer and Zika transcriptomic datasets can be contrasted with RP mutants to determine the accuracy of these hypotheses (i.e. Minute and cancer similarities are not simply due to, for example, cellular stress response, which would also be shared with viral infections).

After constructing gene co-expression modules and identifying hub genes for each RP mutant, we determined the level of preservation between each RP mutant and cancer or Zika datasets (Figure 5; Materials and Methods). This was achieved via comparison of the *D. melanogaster* (17,559 genes) and *Homo sapiens* (56,212 genes) data sets to identify homologues (8,446 genes; Figure 5b). As a conservative direct comparison, these 8,446 homologous genes were used for subsequent WGCNA analysis to determine whether mutations in RPs induce expression changes resembling those in glioblastoma or breast cancer, or Zika virus-infected neuroblastoma cells.

**Figure 5.**
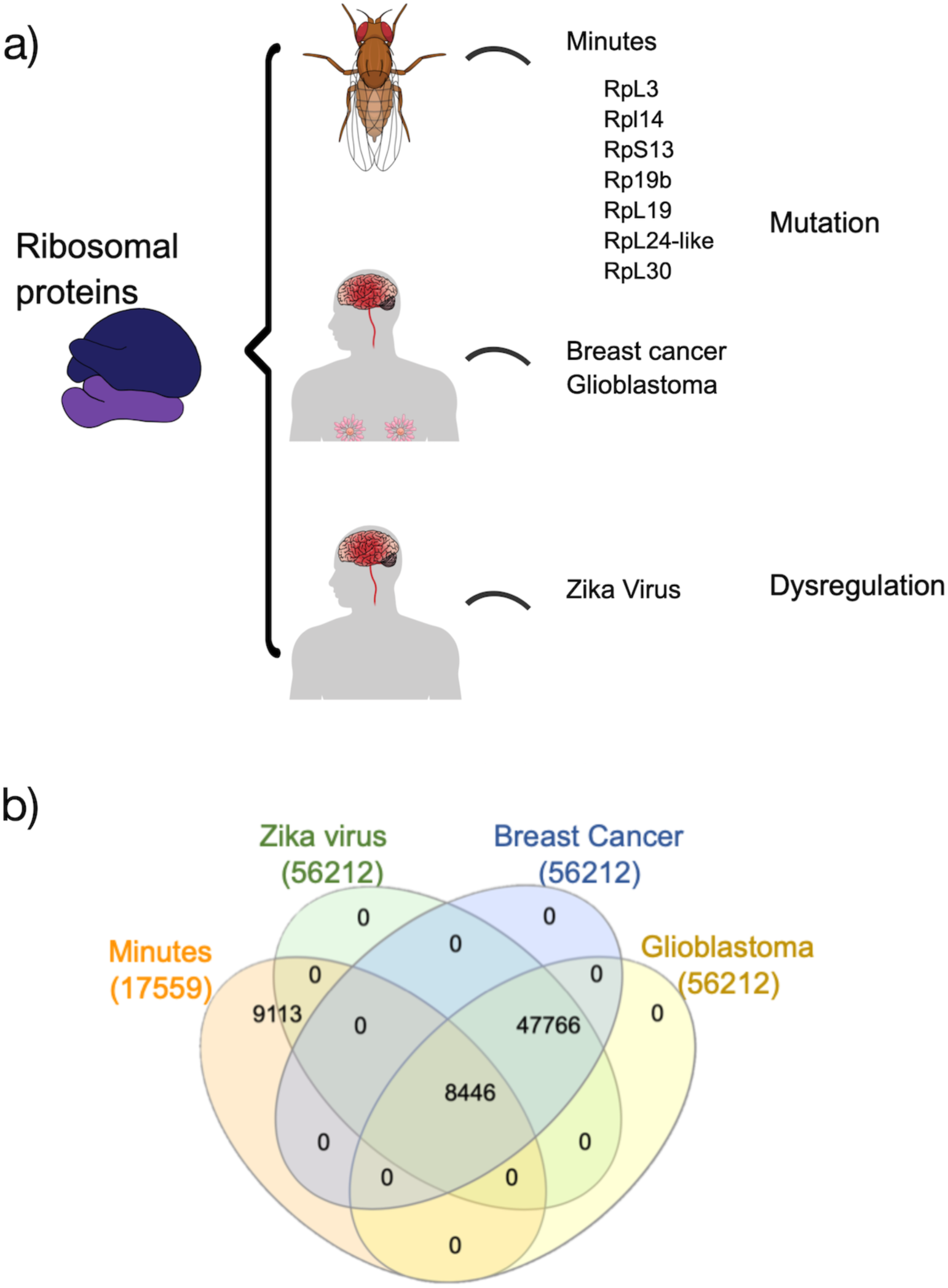
Homologue comparison between fly and human datasets. a) Schematic representation of the WGCNA comparison between Minute mutants and glioblastoma, breast cancer and Zika virus infection transcriptome. b) Venn Diagram of shared genes between Minute mutants and glioblastoma, breast cancer and Zika virus infection.

As in the RP Network Analysis, we employed the modulePreservation function in the WGCNA package, using two network-based composite preservation statistics (Zsummary). We found eight RP modules and 6,729 unique genes having well-defined human counterparts (5 modules and 4,392 genes in glioblastoma and 3 modules and 5,730 genes in breast cancer were highly preserved; Table 6). Like cancer, Zika virus also controls its translation activity through internal ribosomal entry sites, altering translation of ribosomal subunits (Song et al., 2019). Thus, we also compared the preservation of RP hub genes between mutants and the Zika virus infection profile. We did not find any evidence that RP networks are well preserved between mutants and Zika virus infection (Table 6).

**Table 6.** Overview of module preservation statistics. Columns report RP modules and Zika Virus compared and preservation statistics (ZSummary and p-value) for each set of data (Glioblastoma, breast cancer and Zika virus). A Zsummary statistic of >10 is considered strongly similar, >5 moderately similar and <2 as dissimilar.

|  | Glioblastoma vs RPs. |  | Breast Cancer vs RPs. |  | ZikaV vs RPs. |  |
| --- | --- | --- | --- | --- | --- | --- |
|  | Zsummary | p-value | Zsummary | p-value | Zsummary | p-value |
| **RpL3** | <u>9.561</u> | <u><0.0001</u> | <u>7.678</u> | <u><0.0001</u> | 0.094 | 0.710 |
| **RpL14** | <u>10.138</u> | <u><0.0001</u> | -4.550 | 0.977 | -1.425 | 0.929 |
| **RpS13** | <u>7.287</u> | <u><0.0001</u> | <u>5.892</u> | <u>0.0001</u> | 2.178 | 0.149 |
| **RpL19** | -0.739 | 0.8961 | -3.161 | 0.908 | 0.297 | 0.601 |
| **RpS19b** | <u>6.060</u> | <u><0.0001</u> | <u>5.254</u> | 0.0002 | 1.847 | 0.216 |
| **RpL24-like** | <u>12.764</u> | <u><0.0001</u> | <u>9.447</u> | <u><0.0001</u> | 2.242 | 0.116 |
| **RpL30** | <u>8.088</u> | <u><0.0001</u> | <u>4.281</u> | 0.0002 | -0.821 | 0.838 |

We then estimated intramodular connectivity (i.e. connectivity between genes in the same module) of conserved RP modules. Intramodular connectivity was then used to visualise connectedness of RP-hub genes in Minutes (Figure 6). The gene position within the network indicates how it overlaps with other Minutes. Genes occupying a more central position are present in more than one Minute network. Thus, our graph shows that the majority of genes present on the network are shared by more than one RP module, indicating that RP modules are complementary. Furthermore, the network analysis also indicated a high connectedness between different RP genes (i.e. distance between genes inside a cluster) suggesting a common activation pathway.

**Figure 6.**
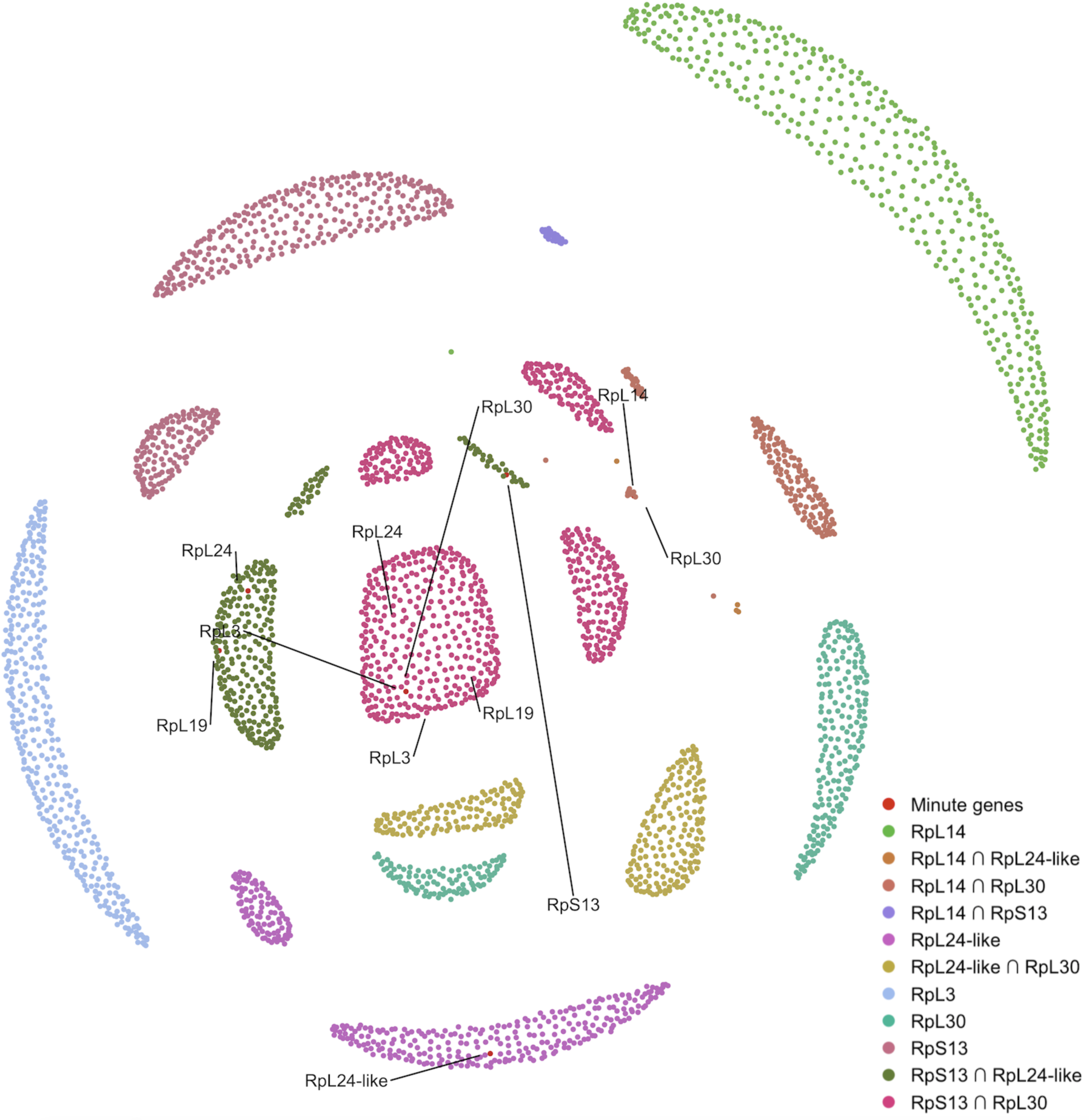
Cluster network of RP genes present in well-conserved modules in glioblastoma data. The co-expression network for each Rp group can be visualised as network components of genes that are exclusive or shared with other modules. Nodes are sized by intramolecular connectivity and coloured by Rp group or shared Rp network. Mutant genes are highlighted in red and labeled.

## Discussion

Here we demonstrate that mutations affecting levels of RPs and causing the Minute phenotype result in highly pleiotropic gene-level transcriptomic changes. When viewed at the level of single-gene differential expression, these lack a clearly defined core of differentially expressed genes across the whole organism. Moreover, there is also no clear pattern or master regulator shaping the expression of RPs. Despite this pleiotropy, when network-based analysis is employed, a high degree of preservation between transcriptomes of all RP mutants was identified. Thus, what may appear stochastic or pleiotropic at one level, reveals a consistent pattern at a different level. This shaping of the entire RP mutant transcriptome bears striking similarity to the transcriptomes of cancer cells.

### Genomic Clustering

We found no evidence of genomic clustering of RP-encoding genes. Gene families are often clustered into “neighborhoods” to facilitate co-regulation (Cera et al., 2019; Spellman and Rubin, 2002). However, this is not the case with *D. melanogaster* RPs, as they appear randomly distributed throughout the genome. Furthermore, previous work identified several RP-encoding genes in close proximity, but did not offer any evidence of co-regulation (Marygold et al., 2007). Co-regulation through genomic clustering is linked to the influence of regulatory elements, such as enhancers, and to chromosome structure. Although effects of these elements can be mitigated by insulating genomic regions, genetic distance may be the most straightforward way to ensure independent regulation. Genomic spacing of RPs potentially permits single-gene transcriptional regulation and greater degree of fine-tuning compensatory regulation as well as promoting a role in monitoring genome stability (Warner and McIntosh, 2009).

### Differential Gene Expression & Regulation

These seven RP mutations lack a clear core of differentially expressed genes, with only 18 being shared among the seven analysed mutants (Supplementary Table S1). Thus, despite tightly controlled regulation and a shared role in ribosome biogenesis, RP mutations alter expression of individual genes in vastly different ways.

Although lacking a core, there remains a high degree of overlap, either in pairwise or sub-group comparisons (Figure 2) between differentially expressed genes in these mutants. The largest overlap of differentially expressed transcripts (441) occurs between RpL24-like, RpS19b, RpL3, and RpL30 (Figure 2). This overlap does not relate to the size of the RP, nor to their position within an assembled ribosome (Anger et al., 2013). The overlap is likely because these mutants have the largest number of differentially expressed transcripts. This certainly does not rule out an underlying biological cause for high levels of differential expression and overlap, but it does reinforce the absence of a clearly defined core.

Finally, and further highlighting variability between their transcriptomes, each of these RPs mutants has a considerable number of uniquely differentially expressed genes (Table 2). These range from 9% of all differentially expressed genes in RpL30 to 28% in RpS13 with an average of 17% across all seven.

It is important to reiterate that our observations are based on four replicates of whole individual flies rather than on pooled samples of specific tissues. Therefore, we would not detect small tissue-specific or temporally-specific responses, but rather macro responses. Although cell-and timing-specific RP functions are critical, they are likely tightly regulated and impossible to generalise. We were unable to detect previously identified *D. melanogaster* RP regulators in either differential expression analysis 2 or Differential Network Analysis (Table 3). This is unsurprising as individual RPs can have highly specific extra-ribosomal functions (Warner and McIntosh, 2009) and are differentially assembled into ribosomes depending on functional or tissue-specific factors (Genuth and Barna, 2018). Rapidly dividing tissues, such as the imaginal disks, likely have elevated or specialised ribosome biogenesis, a conclusion that is supported by the elevated expression of RP encoding genes in this tissue (Brown et al., 2014; Celniker et al., 2009).

In addition, we also examined single-gene markers that have been previously identified as being misregulated in response to mutations affecting RPs (Supplementary Table S2). Once again, no clear core pathway can be identified. Only five genes, asp (cytoskeleton), CycB3 (cell cycle), pik92e (cell and tissue growth), ask1 (JNK pathway) & bsk (JNK pathway), of the twenty candidates examined, representing major cellular regulatory pathways, were misregulated in four or more of the RP mutations. This does not suggest that these cellular processes are not intricately linked to RPs. For example, it is well known that the Toll-apoptosis results in cell elimination of RP mutant cells, when in competition with wild type cells, (Meyer et al., 2014). However, this, like other regulatory or pathway examples, is likely highly context-specific; thus, it is potentially missed when examining RPs at the macro level.

### Dosage Effects

Dosage changes in RP expression, regardless of direction, appear to have similar transcriptomic effects. The strongly overexpressed mutant *RpS19b* resulted in the second highest number of differentially expressed genes (Table 2). Moreover, there was considerable overlap in differentially expressed genes of RpS19b with those of other RP mutants (Figure 2). Dose-dependency is critical in ribosome biogenesis, with overexpression of RPs being linked to disruption of this process (Tye et al., 2019). Moreover, overexpression of several RPs has been linked to cancer (discussed below).

As only one of the seven Minute mutants screened over-expressed the corresponding RP, it is possible that RP overexpression doesn’t generally result in Minute phenotypes. Moreover, the large overexpression of *RpS19b*, compared with underexpression seen with other RPs perhaps suggests that RPs are less sensitive to overexpression. Although this observation does highlight the delicate balance that RPs maintain, further conclusions are impossible without generating a series of strains with overexpressed RP encoding genes

### Function & Network Analysis

The discovery of regulatory networks and pathways are an important step in understanding the mechanisms of complex diseases, and discovering biomarkers and drug targets. As demonstrated in our results, traditional differential gene expression analysis cannot fully grasp the complexity of RP-induced transcriptomic changes. Using two different network analyses (WGCNA and DCE), we identified ribosomal pathways (modules) and how these pathways change in response to ribosomal protein mutations, indicating a large compensatory mechanism. Although both analyses yielded a similar result, we identified RP-hub genes associated with each mutant using WGCNA due to its powerful network estimation. We also showed that these RP modules are highly preserved across all RP mutants strains, as well as displaying a high connectedness (Figure 6). These results indicate the presence of an underlying co-regulation common to all mutants. It also provides strong evidence that RP hub genes are involved in the same biological pathways, as they have similar expression levels and preservation. GO term analysis failed to identify the same pattern, attributing different functions to our RP hub genes and contradicting the network analysis.

GO analysis, while useful, provides only a crude view of how the transcriptomic network is altered. As it relies on a list of differentially expressed genes, a conservative analysis can fail to identify key GO terms. Moreover, differentially expressed genes can fall into several GO biological process categories. Complex traits, such as disease or disruption of key biological processes can result in high numbers of genome-wide associations. In these instances, GO analysis typically correlates with the number of gene members in a specific GO category (Boyle et al., 2017). Therefore, the use of network analysis to group functionally related genes provides a more comprehensive reference framework for the understanding of regulation of ribosomal genes and the networks associated with them.

Given the importance of ribosomal biogenesis for cellular health, growth and homeostasis, it has been proposed that all ribosomal components are tightly co-regulated (Coléno-Costes et al., 2012; Reja et al., 2015; Li et al., 2018). The analysis employed here provides a novel demonstration of interrelated regulation of the RPs.

### Cancer, Zika virus & Ribosomal Proteins

There are several excellent studies that highlight the contributions of RPs to oncogenesis when mutated, and their aberrant expression is often observed in cancerous cells (Bee et al., 2011; Henry et al., 1993; Hong et al., 2014; Lai et al., 2009; Oršolić et al., 2020). In this study, we used WGCNA to construct a co-expression network for identification of gene co-expression modules associated with glioblastoma and breast cancer. We identified 5 RP modules and over 4000 genes closely related to glioblastoma and 3 modules with over 5000 genes related to Breast cancer. Cancers have been widely reported to vary greatly at the transcriptomic level. In fact, different types of cancer arising from the same lineage exhibit tissue-specific gene over-expression (Axelsen et al., 2007). Given the enormous variation in cancer transcriptomics and the RPs analysed, it is reasonable that a target gene approach resulted in a large gene network associated with cancer development.

Disruption of RPs closely mimics transcriptional networks observed in cancers. As one might expect, genes strongly upregulated in several cancers, such as RpL19 (Henry et al., 1993; Hong et al., 2014), do not mimic cancer networks when mutated. Moreover, downregulation of RpL19 in a prostate cancer cell line reduced aberrant growth (Bee et al., 2011). Both differential gene expression and network expression analyses revealed cancer-like transcriptomic responses in RpL24-like and RpS3 mutants. Moreover, all of the Minute RP mutants had at least some cancer markers differentially expressed, or displayed cancer-like transcriptomic networks, especially in RpL24-like mutants.

However, we failed to identify a key common core between RP mutations and Zika virus responses. Flaviviruses, such as Zika virus, have a single-stranded RNA genome that serves as both a messenger for translation and a template for replication (Garcia-Blanco et al., 2016). To efficiently replicate, they depend on cellular translational machinery to produce viral proteins required for replication. As a result, they have evolved efficient mechanisms to modify or even inhibit host protein translation (Sanford et al., 2019). This alteration of host translational mechanisms, as opposed to disruption or overexpression of ribosomal genes, as occurs in RP mutations, could explain why we failed to identify a well-conserved gene core between them. Furthermore, a ModulePreservation comparison between glioblastoma and Zika virus-infected neuroblastoma cells identified a smaller set of preserved RP hub genes than Minutes and glioblastoma, indicating that glioblastoma has a co-expression pattern more similar to RP mutations in fruit flies than to neuroblastoma, further reinforcing the role of RP regulation in cell machinery.

The seven mutant RPs we examined lacked a cancer-like universal marker. Although some RPs misexpressed genetic cancer markers, this was not universal. Of the markers tested, no gene was aberrantly expressed in more than five of the seven RP mutant backgrounds (Table S2).

Our work highlights the potential for *D. melanogaster* Minute mutations to act, not only as models of RPs, but of numerous pathologies. Although there is no shortage of model systems for disease studies, the tools available to *Drosophila* specialists continue to ensure the relevance of this model organism. For example, this work highlights the potential of the Minute system for studying the functions that perturbations in ribosome biogenesis serve in human disease. Moreover, there may also be utility in the system for rapid screening of genes involved in oncogenesis.

### Genome Stability

As RPs assemble in precise relationships into functional ribosomes, imbalances in RP abundance can signal genomic destabilization, which is often associated with cancer development (Penzo et al., 2019). Thus, RPs can have extra-ribosomal roles as sentinels of genome stability (Warner and McIntosh, 2009) and may induce cell cycle arrest and cell death. This suggests that for cells to survive and proliferate in Minute individuals, some regulatory checkpoints have to be overcome in a manner resembling that of precancerous cells.

As shown here, small changes in RP encoding of transcript abundance can have a strong ripple effect throughout the entire transcriptome. Due to their role as gatekeepers of genomic integrity, there is potential for RPs to contribute to hybrid incompatibility. Small differences in regulation of RP-encoding genes between allospecific parents may result in negative fitness outcomes or Minute-like phenotypes. In addition to functions of RPs in oncogenesis and genome integrity, further study of their roles in hybrid incompatibility or underdominance is warranted.

## Conclusion

Although RPs have typically been studied strictly with respect to their roles in ribosome biogenesis, a wealth of research suggests that RPs regulate critical cellular processes. Thus, additional links to cancer are to be expected. In our work, and throughout the literature, RP mutants display non-overlapping pleiotropic phenotypes. Despite the shared Minute phenotype, transcriptomic responses of RP mutants examined here vary greatly at the individual gene level. However, when stepping back and examining not just the trees but the whole forest, it becomes clear that disruption of RPs shapes the transcriptome in similar ways. Furthermore, RP mutations induce changes in *D. melanogaster* that despite the huge evolutionary distances involved, are very similar to those in human cancers.

## Materials & Methods

### Fly Husbandry & Generation of RP Lines

Flies were raised on standard Bloomington media at 24 °C under a 16h light / 8h dark cycle in 50 mL vials.

Several Minute mutants constructed in prior studies by *P*-element insertion mutagenesis were obtained from the Bloomington *Drosophila* Stock Centre (BDSC) and were backcrossed for eight generations to control white-eyed line. Minute males marked with mini-*white w*[-]/Y; P{m*w*}/balancer were crossed to *w*[1] females. mini-*white* female offspring were selected without the balancer and crossed to *w*[1] males. This brought the Minute disruption into a common *w*[1] maternal (mitochondrial and cytoplasmic) and paternal (Y-chromosome) genetic background. Four additional generations of crosses were made in the *w*[1] background before maintaining the stocks as heterozygotes by selecting for m*w*. m*w* flies were crossed to *w*[1] in the final generation before collecting female offspring for RNA extraction. *w*[+] controls (Canton-S) were also generated by crossing wildtype males to w[1] females, selecting *w*[+]/*w*[1] female offspring to cross to *w*[1] males, and backcrossing to w[1] for three additional generations.

During back-crossing of both RpS24 and RpL26 two shades of red segregated flies. This suggests that these strains may contain multiple p-element insertions, and given the low frequency of the second eye colour in subsequent crosses, these p-elements are likely tightly linked. Therefore, these two mutants were removed from the experiment.

### RNA Extraction & Library Preparation

Whole RNA was extracted using a Qiagen RNeasy Micro kit from individual 2-day-old virgin female flies. Biological replicates for each strain listed in Table 7 were performed in quadruplicate (with the exception of RpS24 and RpL26 which have been omitted). RNA quality was assessed using both a NanoDrop (Thermo Scientific), and subsequently a BioAnalyser (Agilent). RNA library construction and subsequent sequencing were performed by the Okinawa Institute of Science and Technology DNA Sequencing Centre – Onna, Okinawa. Libraries were prepared with a TruSeq RNA Library Prep Kit version 2 (Illumina RS-122) and sequenced on an Illumina HiSeq4000 in paired-end mode (PE150).

**Table 7.**
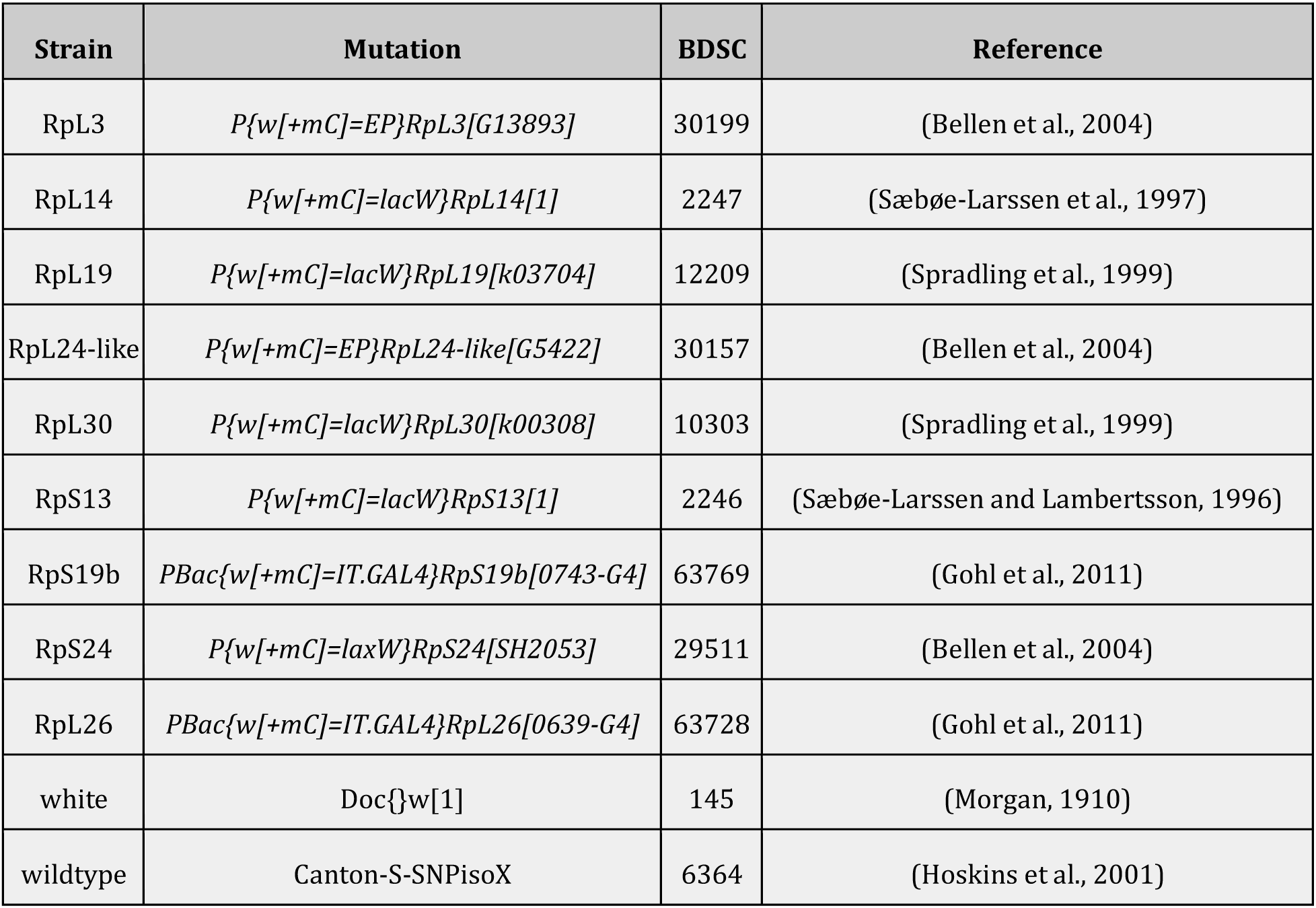
*Drosophila melanogaster* Strains Used.

### Gene Differential Expression Analysis

All statistical analysis and results can be viewed at https://github.com/marivelasque/Minute_project.git. RNA data were trimmed with Trimmomatic 0.38 (Bolger et al., 2014) and quality was assessed using FASTQC (Andrews, 2010). Transcript abundance was calculated with RSEM/bowtie2 (Langmead and Salzberg, 2012; Li and Dewey, 2011). Mapping and abundance calculations were performed against the *D. melanogaster* genome assembly BDGP6 (release 89). Differential expression analysis was performed using DESEQ2 (Love et al., 2014). Visualization relied on packages UpSetR (Conway et al., 2017), ggplot2 (Wickham, 2009, p. 2), igraph (Csardi and Nepusz, 2006), circlize (Gu, 2014) and chorddiag (Gu, 2014). Analysis was run and figures generated using R in the RStudio environment. A markdown file containing the analysis pipeline and generated figures is available on GitHub (updated prior to publication).

### Data Filtering

For both the differential gene expression analysis, conducted with DESEQ2 and WGCNA analysis, genes differentially expressed between white and wildtype were removed. A total of 1332 genes were differentially expressed between the white and wildtype. Removal of these genes from the analysis serves as a conservative control for the presence of a functional/semi-functional copy of the *white* gene.

### Network Analysis

To estimate Differential Network Expression, we used the dnapath R package. To control the network differences in white, we removed all genes significantly co-expressed between white and wildtype. We removed a total of 1750 genes that were differentially co-expressed in white. Differential Network Expression was estimated using pairwise comparisons.

We used the WGCNA R package to construct the co-expression network using gene counts obtained from RSEM. Prior to the analysis, we removed gene outliers using the “goodSamplesGenes” function and accessed overall sample clustering in the expression matrix using “hclust.” We used the lowest threshold power that resulted in approximate scale-free topology to construct the gene co-expression network using “blockwiseModules.” Then, we estimated the gene expression profile, module eigengene, and the module-treatment relationship, calculating the correlation between the module eigengene and treatment group. Modules that were significantly associated with the treatment were used to generate a gene network plot with the R package, igraph. We merged all treatment groups, generating a gene network plot for all treatment groups (Figure 6). Individual plots (i.e., each treatment group) can be found in the Supplemental Material (Figures S1-6).

To determine whether RP mutant modules were conserved groups, we used a WGCNA integrated function (modulePreservation) to calculate module preservation statistics, applied using two network-based composite preservation statistics (Zsummary). Zsummary results and respective p-values were generated through permutation testing (500 permutation). A Zsummary statistic of >10 is viewed as being strongly similar, >5 moderately similar and <2 as having no similarity (Langfelder et al., 2011). The R script used in this study is available at https://github.com/marivelasque/Minute_project.git.

We examined preservation of network properties of the ribosomal hub genes from mutants and two types of cancer, glioblastoma (Bioproject PRJNA388704) and breast cancer (Bioproject PRJNA484546; Rao et al., 2019) and Zika virus (Bioproject PRJNA497590; Martín et al., 2018). Raw reads were downloaded from NCBI and mapping and abundance calculations were performed using the same pipeline used for Minute data against the Genome Reference Consortium Human Build 38.

To establish universal gene identifiers and to facilitate comparisons between human and Minute data, we re-annotated human gene identifiers with *Drosophila melanogaster* gene orthologs. Only identifiers that were common to both data sets (human and fruit flies) were retained. We used the same approach to construct the gene co-expression network as described above. Next, we compared preservation of RP modules between human and Minute data using modulePreservation statistics from WGCNA software.

Network plots were generated by building an adjacency matrix from the RP group and module. The adjacency matrix was then converted into an igraph object and merged by gene name.

## Declarations

## Acknowledgements

We thank Kevin R. Cook (Indiana University Bloomington) and Steven J. Marygold (University of Cambridge) for advice, suggestions, comments on the manuscript and discussion of Minutes. We thank Heather Flores and Jarek Bryk for comments on the manuscript. We thank Steve D. Aird for editing the manuscript. Stocks obtained from the Bloomington *Drosophila* Stock Center (NIH P40OD018537) were used in this study. The Okinawa Institute of Science & Technology Sequencing Centre performed the RNA library preparations and generated the sequencing data.

## Funding

The experiments and analysis were funded by the University of Hawai’i at Mānoa, National Institutes of Health COBRE P20 GM125508 awarded to FAR, the Okinawa Institute of Science & Technology and JSPS KAKENHI Grants 19K06795 and 19K16205 awarded to JAD and MV respectively. JAD and MV were supported by the Okinawa Institute of Science & Technology Genomics & Regulatory Systems Unit.

## Availability of Data & Materials

All code is available at and a markdown file showing all analysis will be available on GitHub prior to publication (https://github.com/marivelasque/Minute_project.git.). All raw sequencing data will be made available prior to publication under the BioProject PRJDB10781.

## Author Contributions

FAR conceived the project. JAD and FAR conducted the experiments. JAD, MV and FAR analyzed the results and wrote the paper.

## Competing Interests

The authors declare no competing interests.

## Supplementary Figures

**Table S1.** Core Genes. Gene names and IDs for differentially expressed genes (adj-p <0.05) that are common to all seven RP mutant lines. No filtering was performed for the direction of differential expression change.

| Gene | Gene ID |
| --- | --- |
| *Arr2* | *FBgn0000121* |
| *cni* | *FBgn0000339* |
| *CRMP* | *FBgn0023023* |
| *CG3740* | *FBgn0023530* |
| *CG14476* | *FBgn0027588* |
| *CG2930* | *FBgn0028491* |
| *CG1561* | *FBgn0030317* |
| *CG5613* | *FBgn0030839* |
| *CG8560* | *FBgn0035781* |
| *BoYb* | *FBgn0037205* |
| *CG2943* | *FBgn0037530* |
| *mask* | *FBgn0043884* |
| *CG31077* | *FBgn0051077* |
| *Atox1* | *FBgn0052446* |
| *Muc14A* | *FBgn0052580* |
| *Ctr1B* | *FBgn0062412* |
| *AGO2* | *FBgn0087035* |
| *CR43961* | *FBgn0264676* |

**Table S2.**
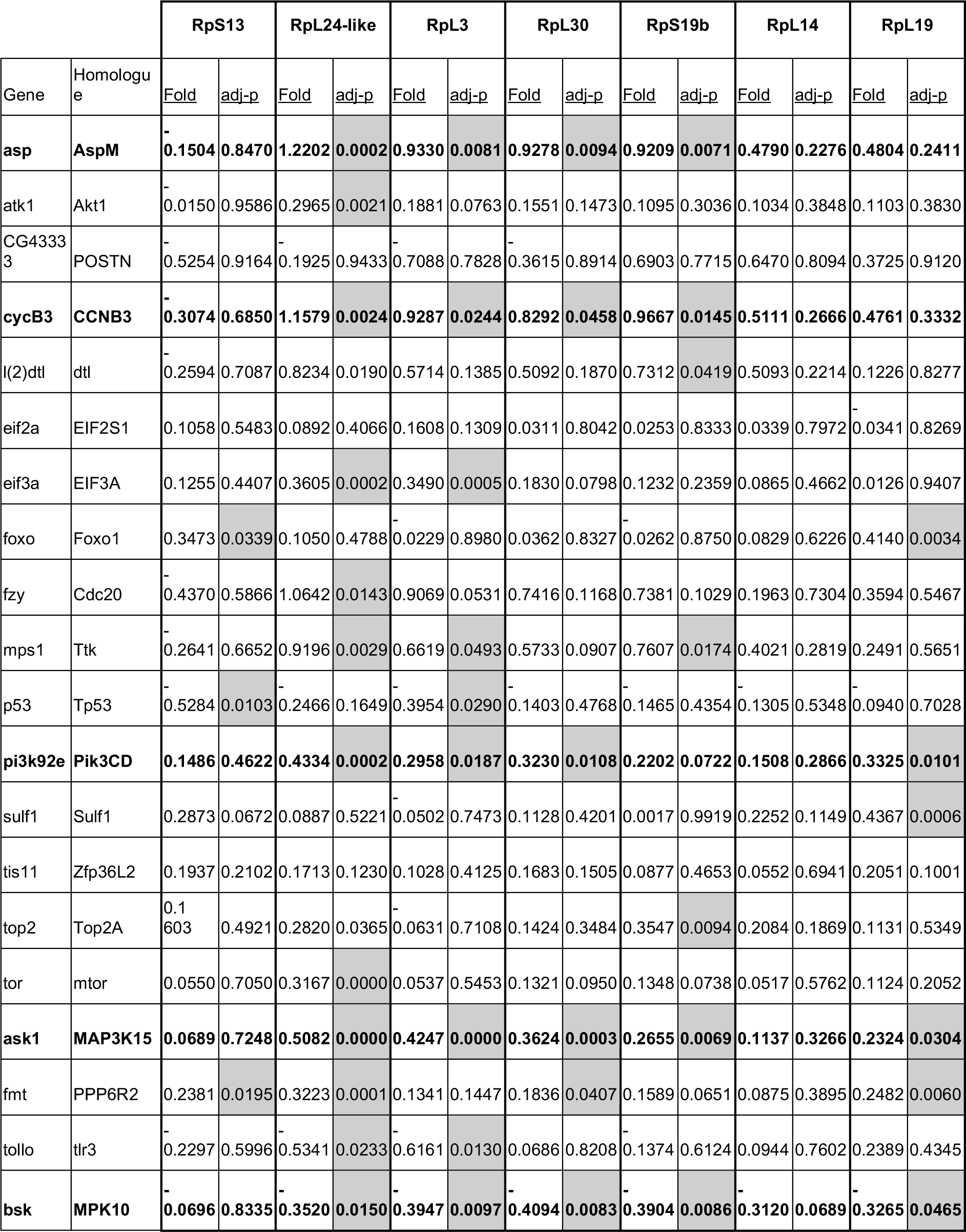
Differential Expression of Candidate Genes.

**Table S3.**
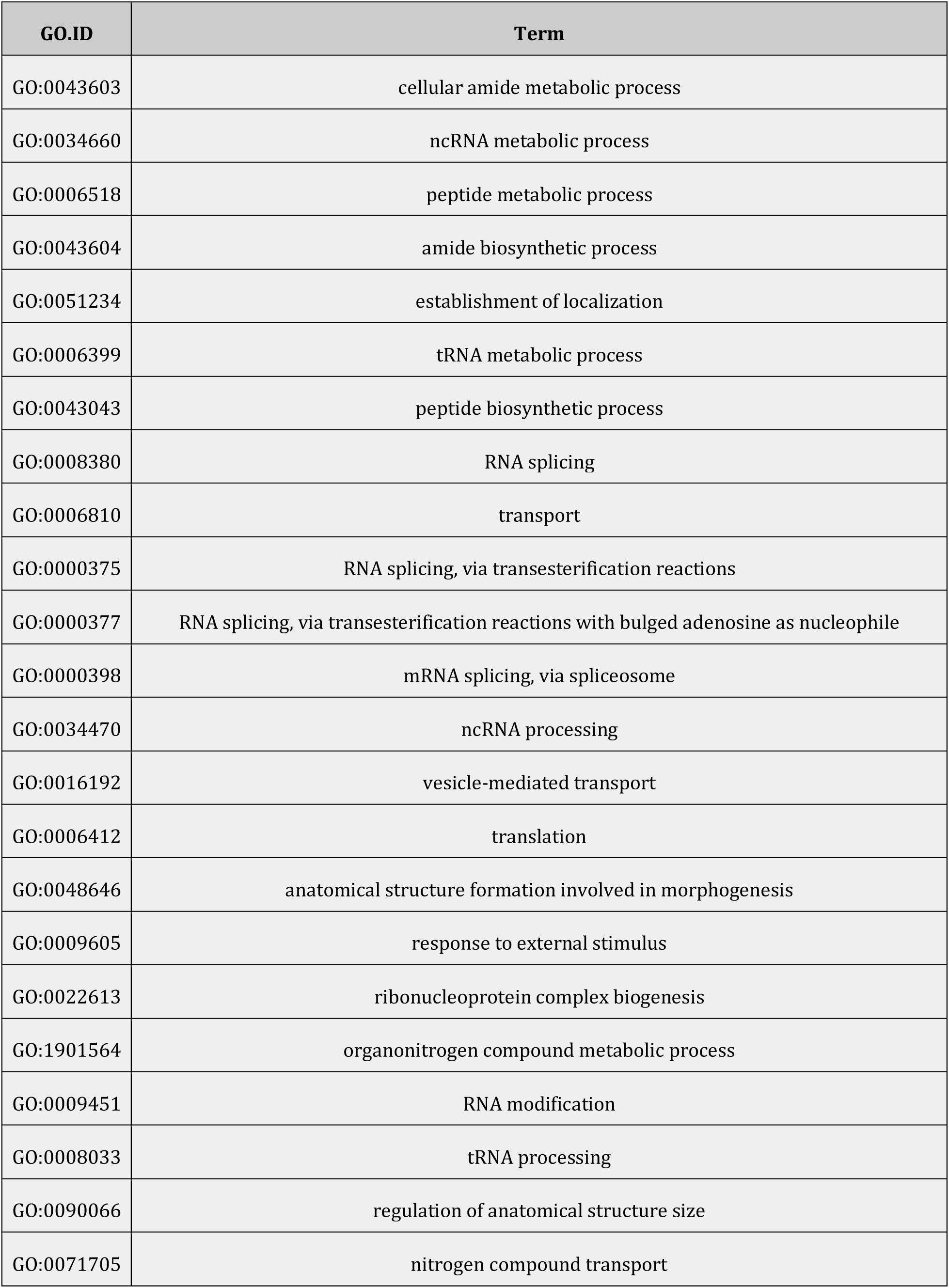
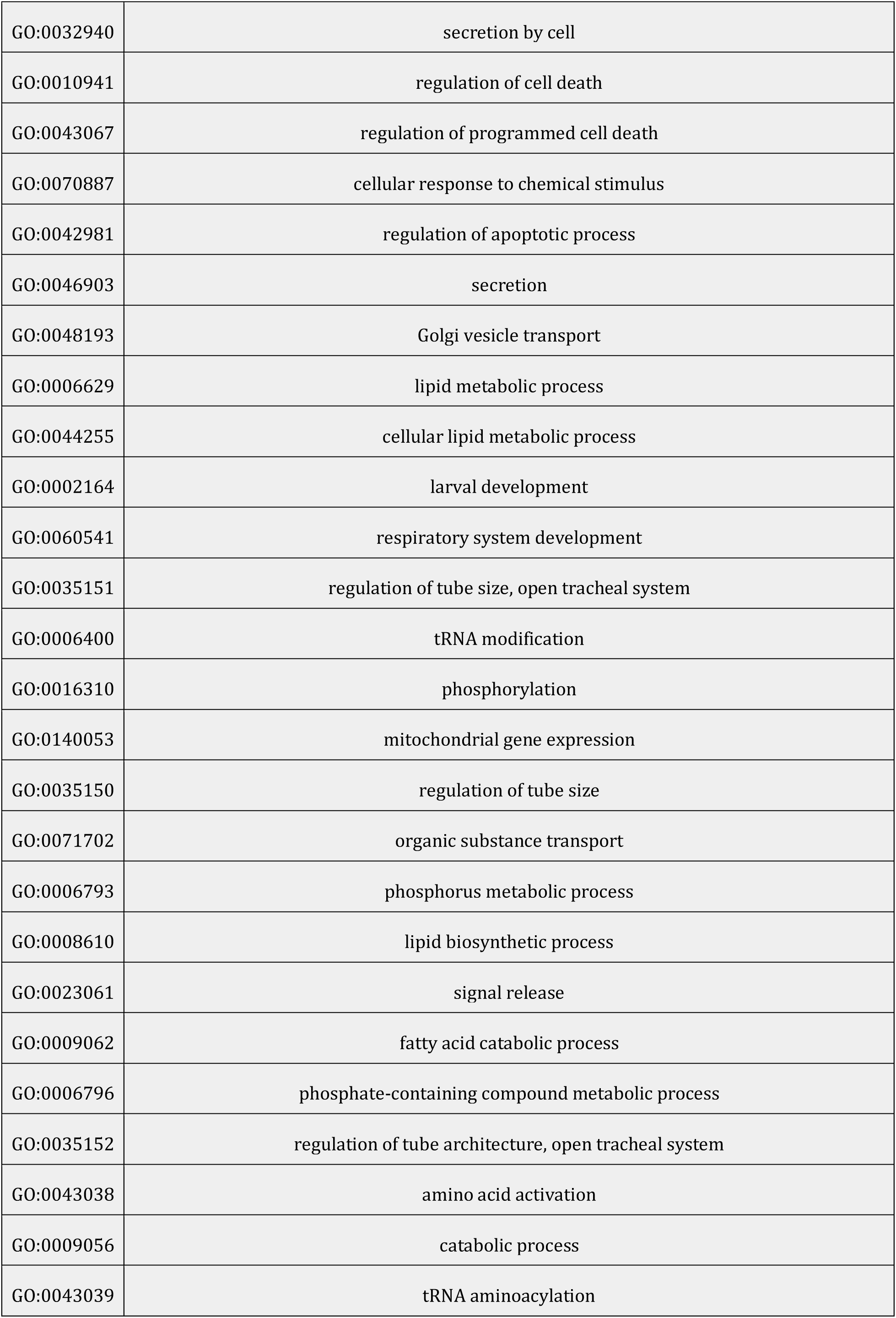
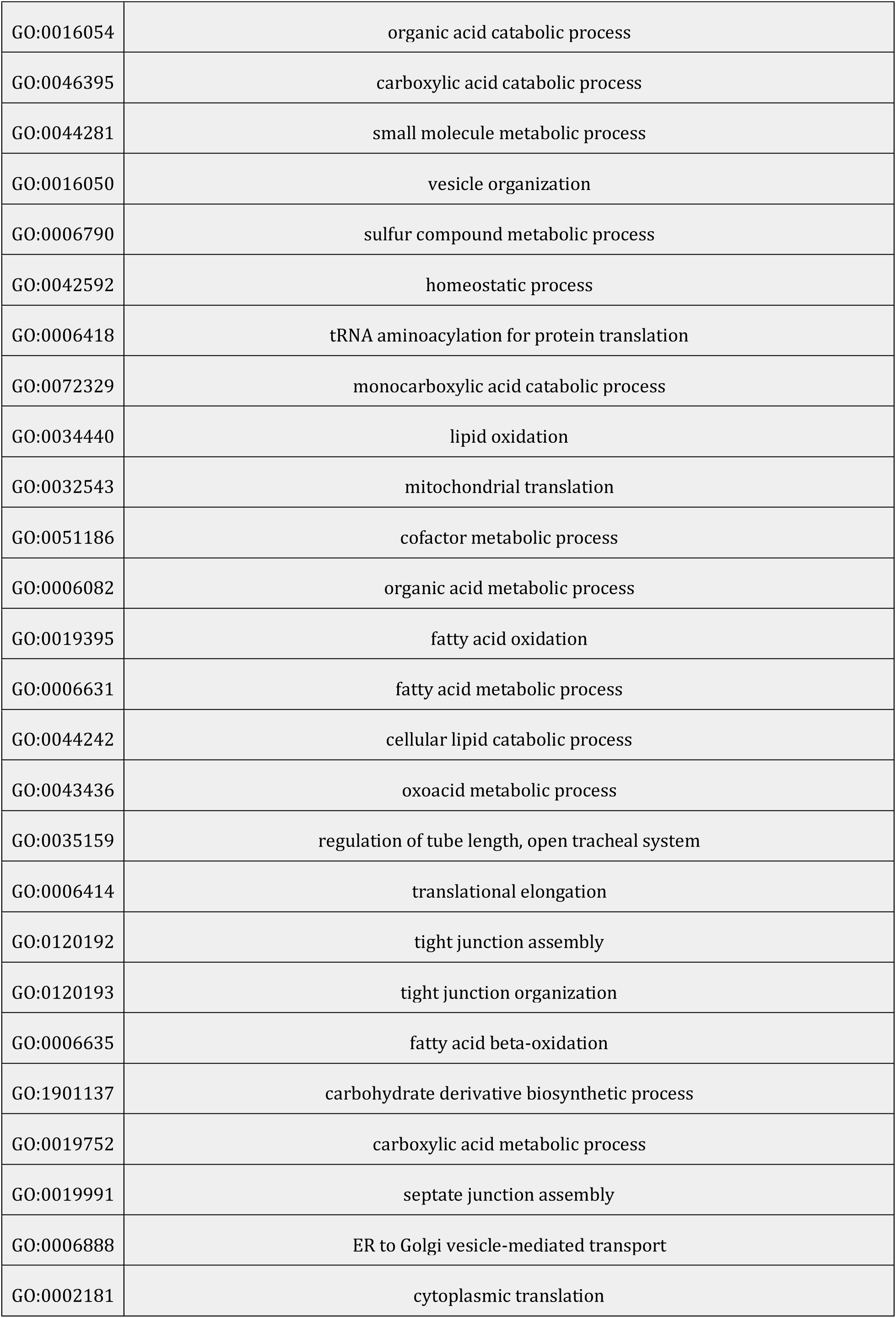
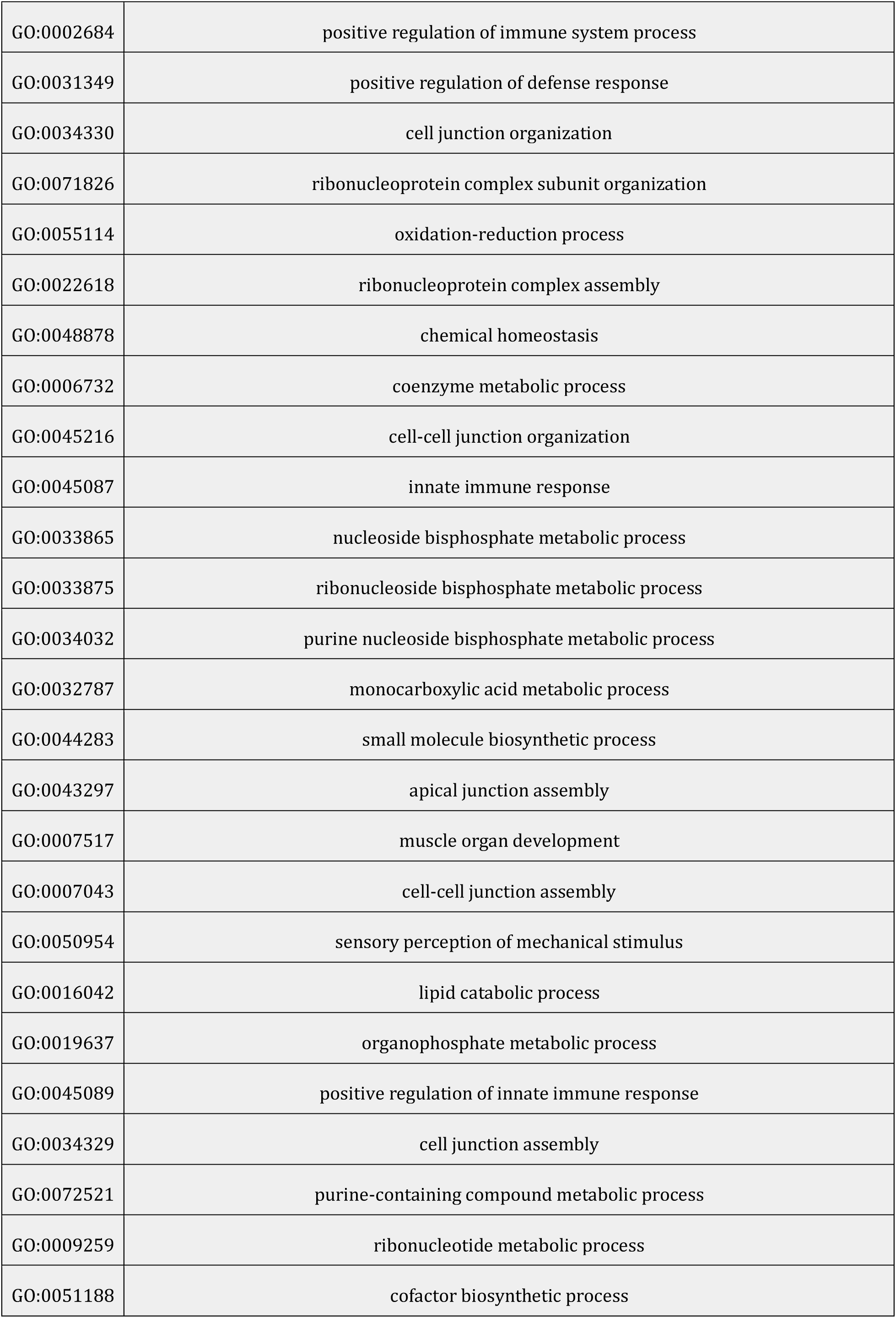
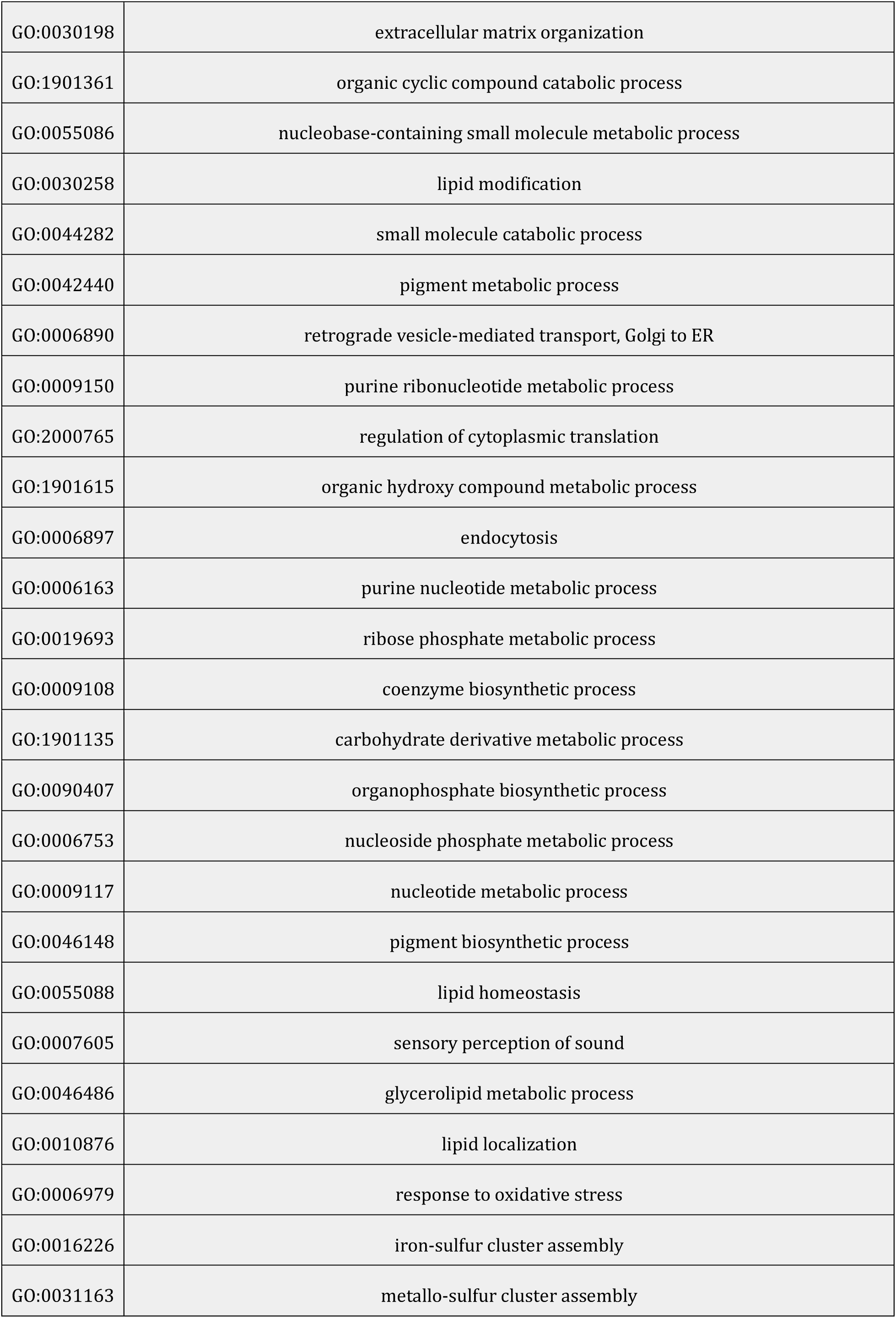
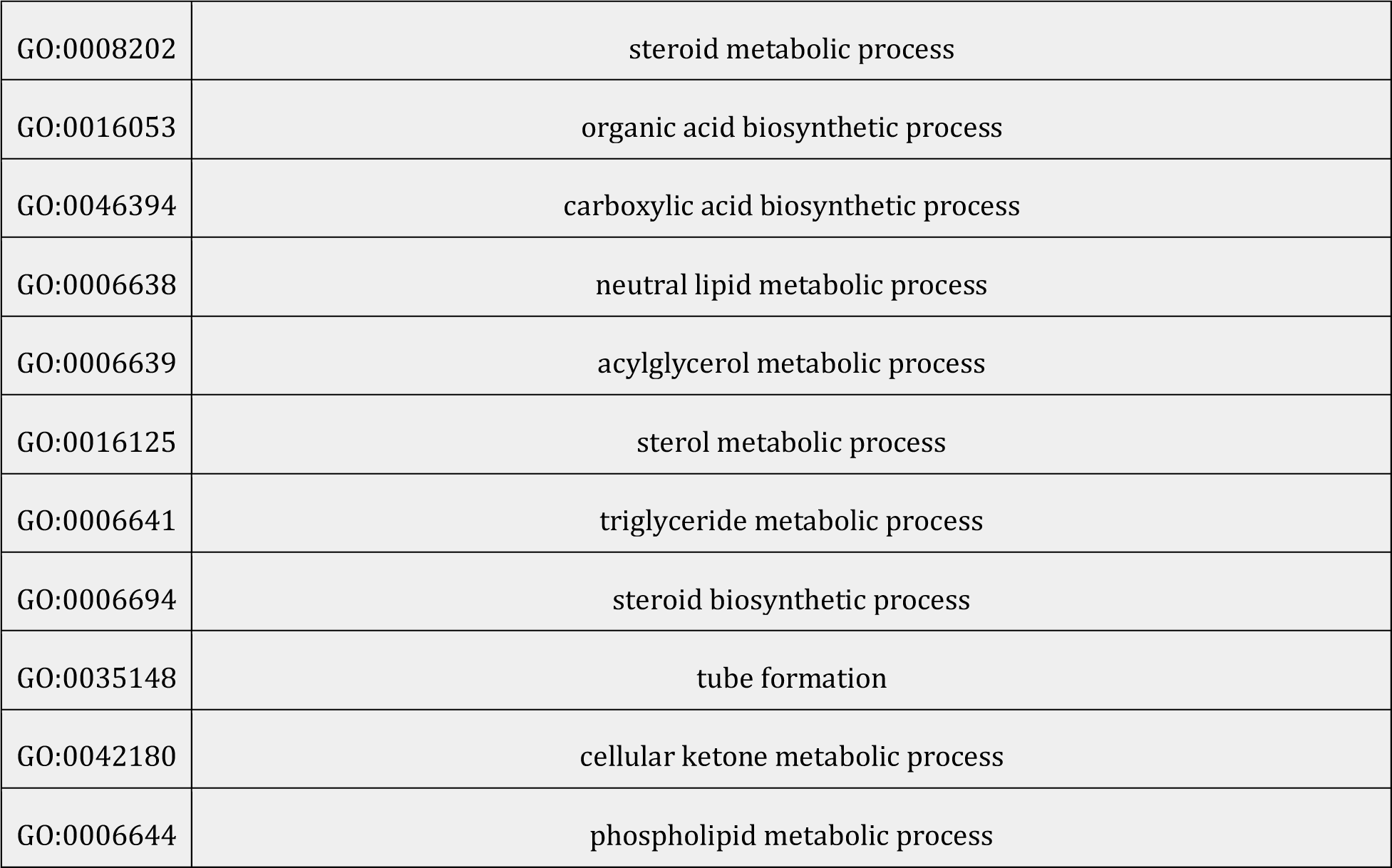
Shared Gene Ontology Categories of RP Mutants.

**Figure S1.**
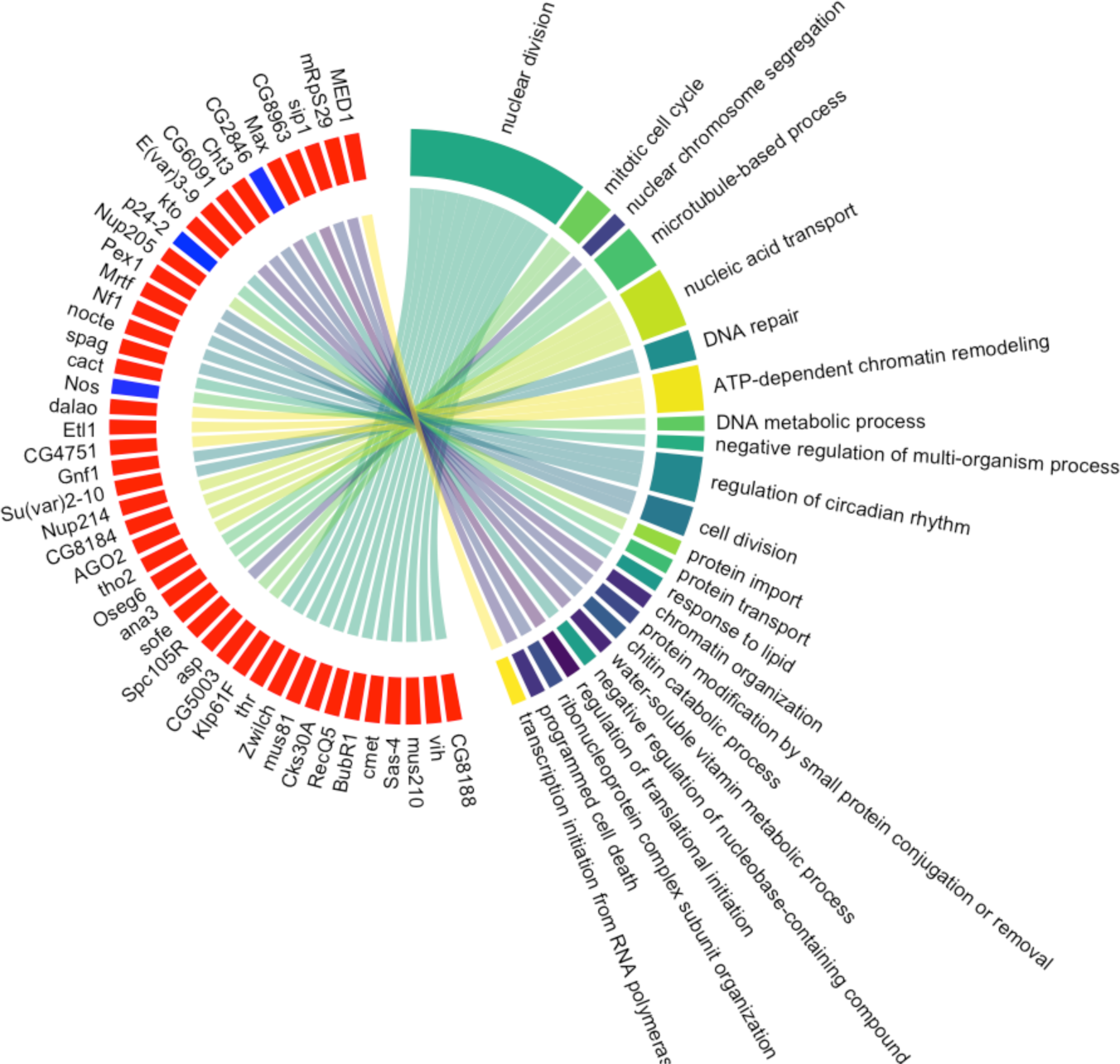
Circle plot of RpL3 mutants indicating the association between differentially expressed genes inside the “RpL3” gene cluster and their associated GO terms. Associations between individual genes and their terms are indicated with ribbons. Genes upregulated in the RpL3 module are represented in red and those downregulated in blue.

**Figure S2.**
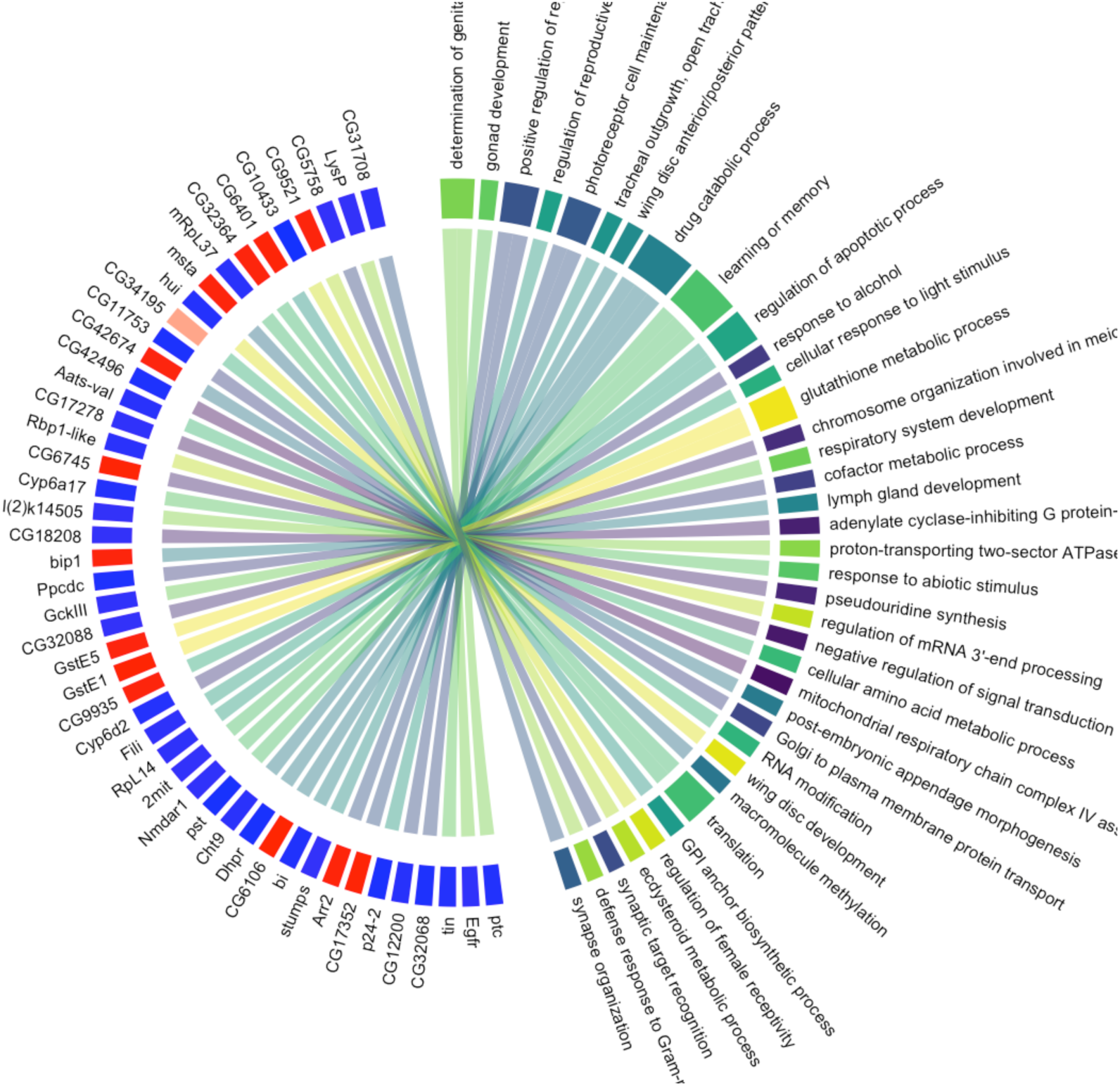
Circle plot of RpL14 mutants indicating the association between differentially expressed genes inside the “RpL14” gene cluster and their associated GO terms. Associations between individual genes and their terms are indicated with ribbons. Genes upregulated in the RpL14 module are represented in red and those downregulated in blue.

**Figure S3.**
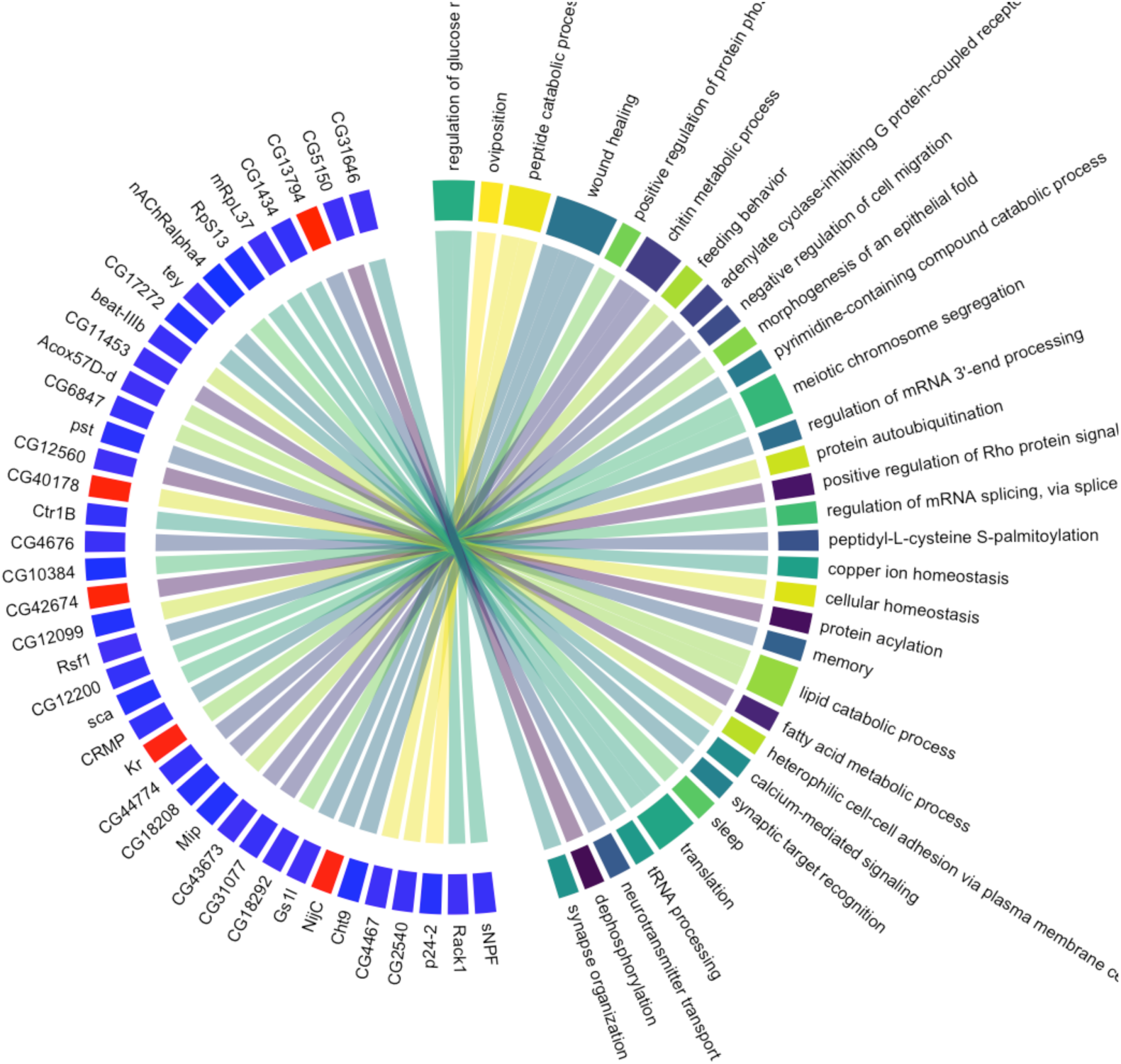
Circle plot of RpS13 mutants indicating the association between differentially expressed genes inside the “RpS13” gene cluster and their associated GO terms. Associations between individual genes and their terms are indicated with ribbons. Genes upregulated in the RpS13 module are represented in red and those downregulated in blue.

**Figure S4.**
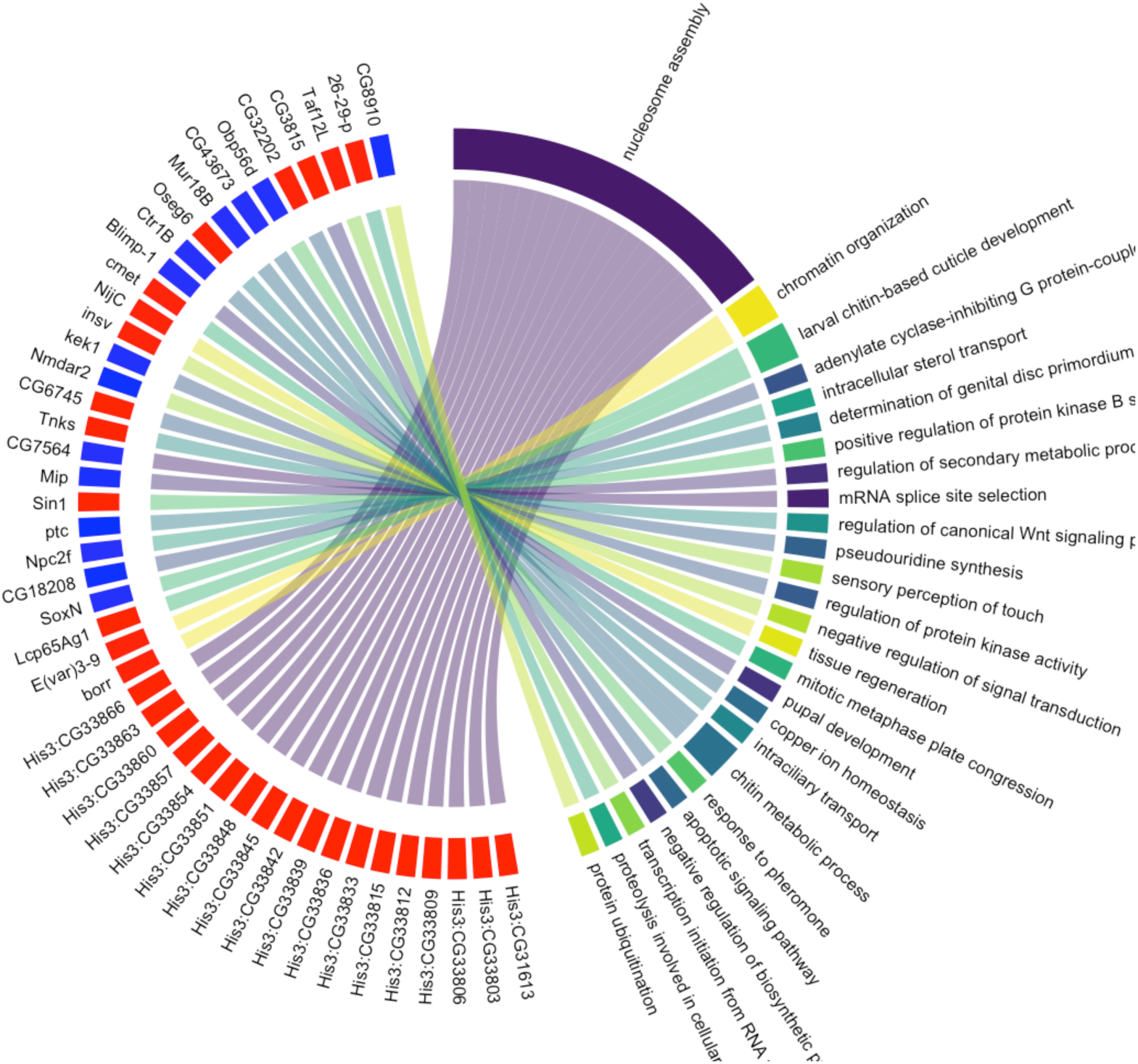
Circle plot of RpS19b mutants indicating the association between differentially expressed genes inside the “RpS19b” gene cluster and their associated GO terms. Associations between individual genes and their terms are indicated with ribbons. Genes upregulated in the RpS19b module are represented in red and those downregulated in blue.

**Figure S5.**
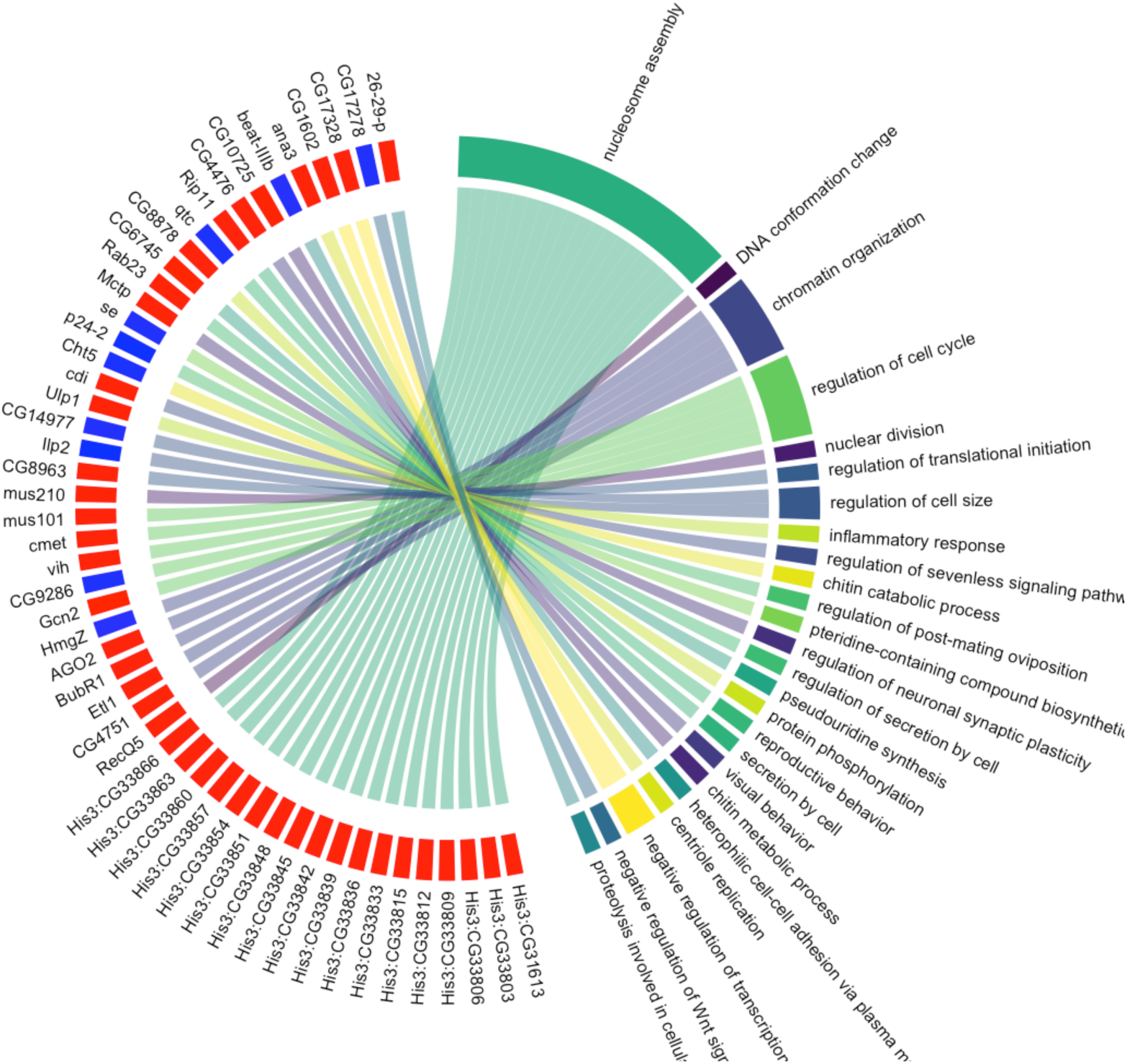
Circle plot of RpL24-like mutants indicating the association between differentially expressed genes inside the “RpL24-like” gene cluster and their associated GO terms. Associations between individual genes and their terms are indicated with ribbons. Genes upregulated in the RpL24-like module are represented in red and those downregulated in blue.

**Figure S6.**
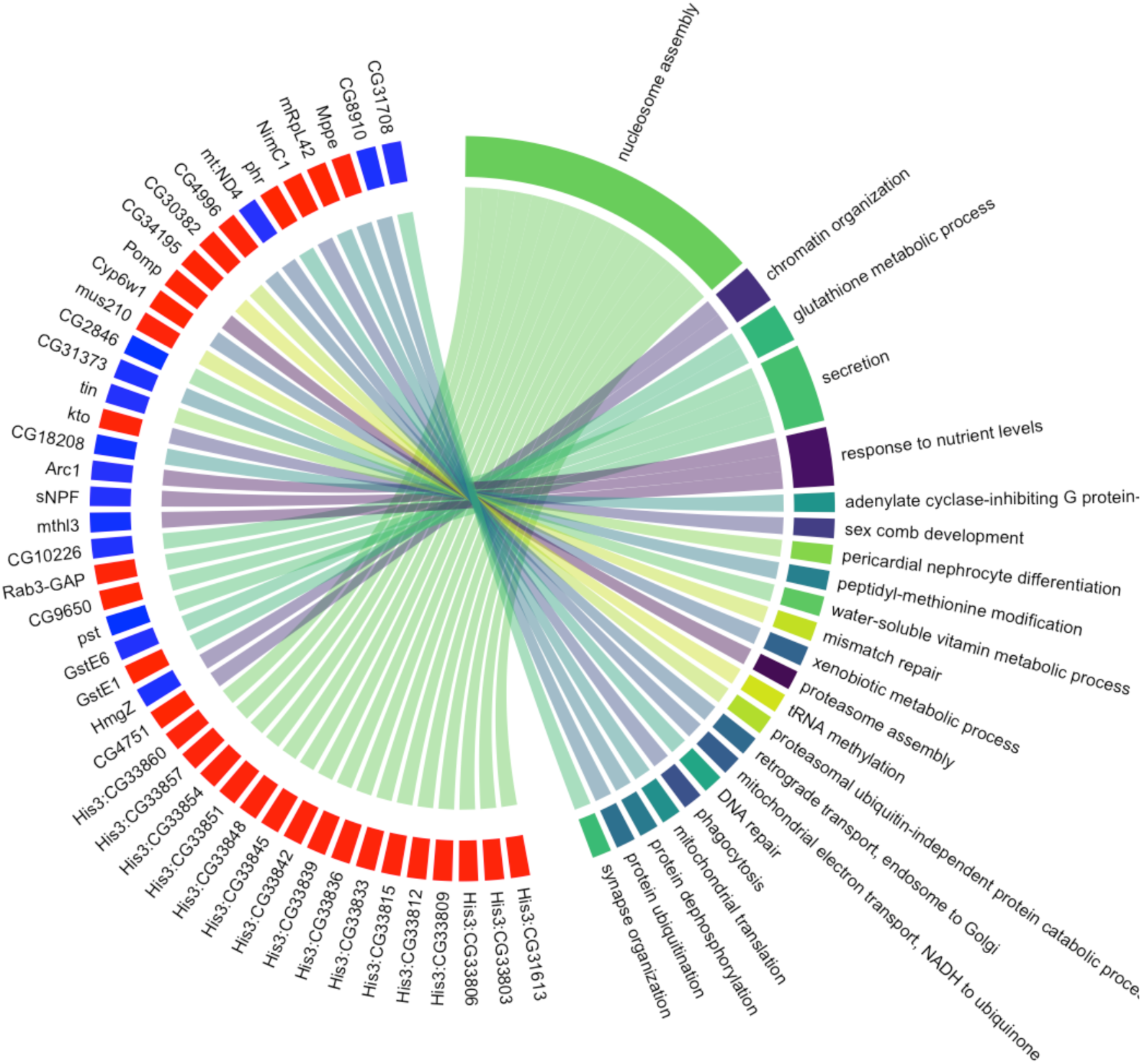
Circle plot of RpL30 mutants indicating the association between differentially expressed genes inside the “RpL30” gene cluster and their associated GO terms. Associations between individual genes and their terms are indicated with ribbons. Genes upregulated in the RpL30 module are represented in red and those downregulated in blue.

**Figure S7.**
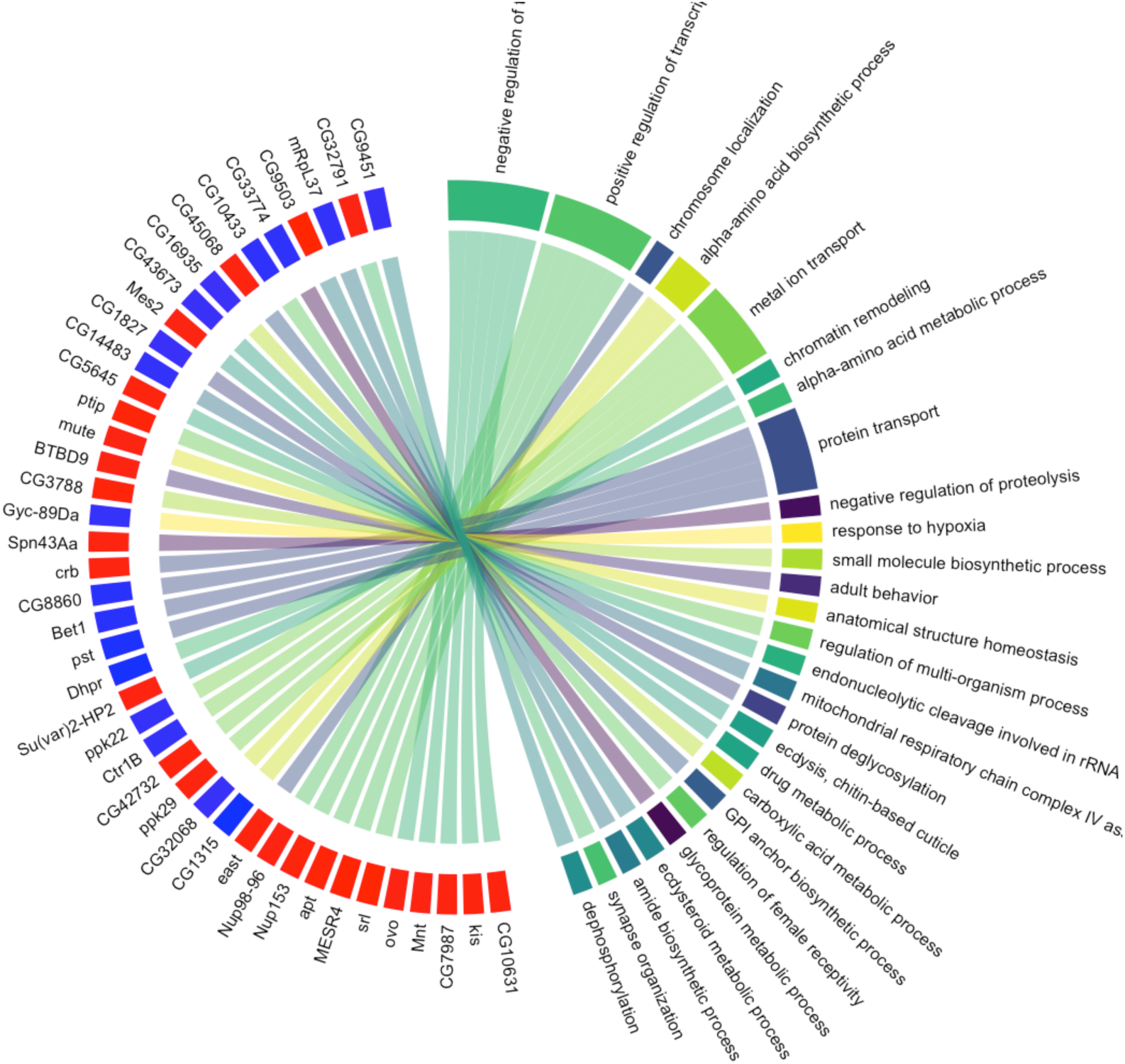
Circle plot of RpL19 mutants indicating the association between differentially expressed genes inside the “RpL19” gene cluster and their associated GO terms. The association between individual genes and their GO terms is indicated via a ribbon. Upregulated genes in the RpL19 module are represented in red and downregulated genes in blue.

## Notes

### Competing Interest Statement

The authors have declared no competing interest.

